# Cryo-plasma FIB/SEM volume imaging of biological specimens

**DOI:** 10.1101/2022.09.21.508877

**Authors:** Maud Dumoux, Thomas Glen, Elaine M. L. Ho, Luís M. A. Perdigão, Sven Klumpe, Neville B.-y. Yee, David Farmer, Jake L. R. Smith, Pui Yiu Audrey Lai, William Bowles, Ron Kelley, Jürgen M. Plitzko, Liang Wu, Mark Basham, Daniel K. Clare, C. Alistair Siebert, Michele C. Darrow, James H. Naismith, Michael Grange

## Abstract

Serial focussed ion beam scanning electron microscopy (FIB/SEM) enables imaging and assessment of sub-cellular structures on the mesoscale (10 nm to 10 μm). When applied to vitrified samples, serial FIB/SEM is also a means to target specific structures in cells and tissues while maintaining constituents’ hydration shells for *in-situ* structural biology downstream. However, the application of serial FIB/SEM imaging of non-stained cryogenic biological samples is limited due to low contrast, curtaining and charging artefacts. We address these challenges using a cryogenic plasma FIB/SEM (cryo-pFIB/SEM). We evaluated the choice of plasma ion source and imaging regimes to produce high quality SEM images of a range of different biological samples. Using an automated workflow we produced three dimensional volumes of bacteria, human cells, and tissue, and calculated estimates for their resolution, typically achieving 20 to 50 nm. Additionally, a tag-free tool is needed to drive the application of *in situ* structural biology towards tissue. The combination of serial FIB/SEM with plasma-based ion sources promises a framework for targeting specific features in bulk-frozen samples (>100 μm) to produce lamella for cryogenic electron tomography.

## Introduction

Volumetric imaging using a dual-beam focused ion beam/scanning electron microscope (FIB/SEM) enables large volumes of material to be reconstructed into 3D contextual maps, for cells and tissues. However, this technique is almost exclusively applied to fixed, resin-embedded, and stained samples. There are only a few studies demonstrating application of this technique to vitrified, frozen-hydrated material (Scher et al., 2021; Schertel et al., 2013; Zhu et al., 2021). Vitrified (as opposed to fixed) samples are required where volumetric information is to be used for further *in-situ* structural studies by cryogenic electron tomography (cryo-ET). Cryo-ET can determine protein structures in the context of a cell at pseudo atomic resolution but requires locating the regions of interest (ROI) within the experimental specimen (known as a lamella) which should be sufficiently thin to be transparent to transmitted electrons (< 300 nm). Lamellae are most fabricated with cryo focused ion beam scanning electron microscopes (cryoFIB/SEM) (Bäuerlein and Baumeister, 2021). Combining ROI targeting and serial FIB/SEM analysis of biological volumes is an attractive approach to multi scale imaging.

The process of thinning (milling) requires a focused ion beam; the current state of the art uses Ga^+^ ion beams. For lamella preparation, ensuring the ROI is contained within the lamella commonly employs correlative light microscopy (cryo-CLEM) relying on fluorescent marker (usually a tagged protein) prior to, and during milling (DeRosier, 2021; Klein et al., 2021; Mahamid et al., 2015). However, fluorescence microscopy under cryogenic conditions suffers from poor axial resolution (> 300 nm), reducing its effectiveness as a targeting tool for lamella preparation. Moreover, the shuttling between machines can introduce alignment / handling errors. As structural biology moves to human tissue, developing approaches that do not depend on fluorescence to identify ROIs will be critical. An idealised approach would identify structural features in real time during milling and thus enable localisation of the ROI within 50 nm (thus ensuring the ROI is contained even in the thinnest lamella).

There are two alternatives to cryogenic fluorescence imaging: cryo-soft-X-ray tomography (Kounatidis et al., 2020) or serial FIB/SEM (Schertel et al., 2013; Spehner et al., 2020). Cryo-soft-X-ray tomography can image volumes up to 10 μm thick at 40 nm resolution or 1 μm thick at 25 nm and has been successfully combined with fluorescence microscopy (Kounatidis et al., 2020). However, X-ray tomography requires a synchrotron and soft X-ray induced beam damage which can adversely affect the sample, damaging higher resolution features and precluding cryo-ET experiments.

Serial FIB/SEM is a well-known volumetric imaging technique: a layer of surface is removed by the FIB and the newly exposed surface is imaged by SEM in a repeated cycle (hence serial). For this to be useful for localisation of ROI for cryo-ET, it needs to be carried out on vitrified samples. Current cryo-FIB/SEM approach can be extended further through the introduction of an inductively coupled plasma FIB (pFIB) generated from gases as an alternative to Ga^+^ ions. Such pFIB ion sources can be operated at higher currents than Ga^+^ (Burnett et al., 2016), thus allowing sufficiently rapid ablation rates that is needed to remove the large volumes of material found in tissue samples (as opposed to single cells). A potential advantage is a reduction in ion implantation into the sample, which is known for Ga^+^ in solid state chemistry (Eder et al., 2021).

Imaging native contrast of biological material using serial FIB/SEM is difficult. The interaction with low atomic number elements results in fewer backscattered and consequently secondary electrons (Reimer and Tollkamp, 1980); therefore, images have low sample contrast (Reimer and Tollkamp, 1980). Sample contrast has traditionally been improved by sample fixation and staining with heavy metal. However, this leads to ultrastructural changes and dehydration, such that constituent proteins cannot be analysed to near atomic resolution. Therefore, increasing the contrast generated from native samples is important to extend its application to a cryo-ET workflow. For insulating materials, such as biological materials, there are two primary electron energies where the secondary electron yield is equal to the number of incident electrons – so called crossover energies. Imaging at these energies stabilises the potential of the surface and reduces charging (Joy and Joy, 1996; Seiler, 1983). Therefore, implementing acquisition schemes that utilise these crossover energies, while imaging with a short working distance electrostatic lens, may lead to greater contrast and suitability of serial FIB/SEM for targeting.

Here we report serial cryogenic plasma FIB/SEM (cryo-Serial pFIB/SEM) analysis of vitrified hydrated biological specimens. We evaluated different plasma gases for their suitability and obtained serially generated volumes with high biological contrast. We describe an automated data collection routine that maximises image contrast and feature localisation. Using this approach we imaged a range of samples from bacteria, cells and tissue demonstrating the ability of the technique applied to vitrified material to generate sub-cellular information that may be suitable for targeting. For photosynthetic bacteria, we were able to observe nascent chromatophores, in mammalian cells we quantified the extent of membrane contact sites and mitochondrial network, in vitrified mouse brain we observed synapses and synaptic contents, while in mouse heart we were able to observe cellular organisation of sarcomeres and other organelles within the tissue.

## Results

### Plasma FIB-characterisation for FIB/SEM on plunge frozen samples

SEM imaging is extremely sensitive to variations in surface topography, so even small distortions (known as curtains) induced by FIB milling give rise to features parallel to the milling direction (appearing as vertical lines on the SEM images). The features obscure the underlying structure and thus their presence should be eliminated. We assessed the degree to which curtaining occurred on vitreous cellular samples while milling with plasma at different currents (Figure 1). We determined the curtaining propensity (as %) at three ion currents (high, medium, and low) to analyse this relationship for four different gases argon, xenon, nitrogen, and oxygen at two different voltages (30kV and 20kV) (Figure 1; Figure 1 - Supplementary Figure 1 to 3). Representative images and associated current are shown in Figure 1 and Figure 1 Supplementary Figure 2 and 3. Variations in the curtaining could be determined from the data, with a trend towards greater curtaining propensity exhibited at high currents vs low currents. The notable exception to this is nitrogen at 20 kV (Figure 1 - supplementary figure 2). Nitrogen and oxygen plasmas exhibit the greatest proportion of curtaining artefacts compared to xenon and argon both at high, medium, and low currents. This suggests that plasmas generated from xenon and argon produce the smoothest surfaces for bulk milling and pFIB/SEM imaging. (Figure 1). Both argon and xenon have a curtaining propensity under 10% at low (around 0.2 nA) pFIB/SEM imaging currents. However, argon presents lesser propensity for curtaining at greater currents than xenon during bulk milling.

**Figure 1.**
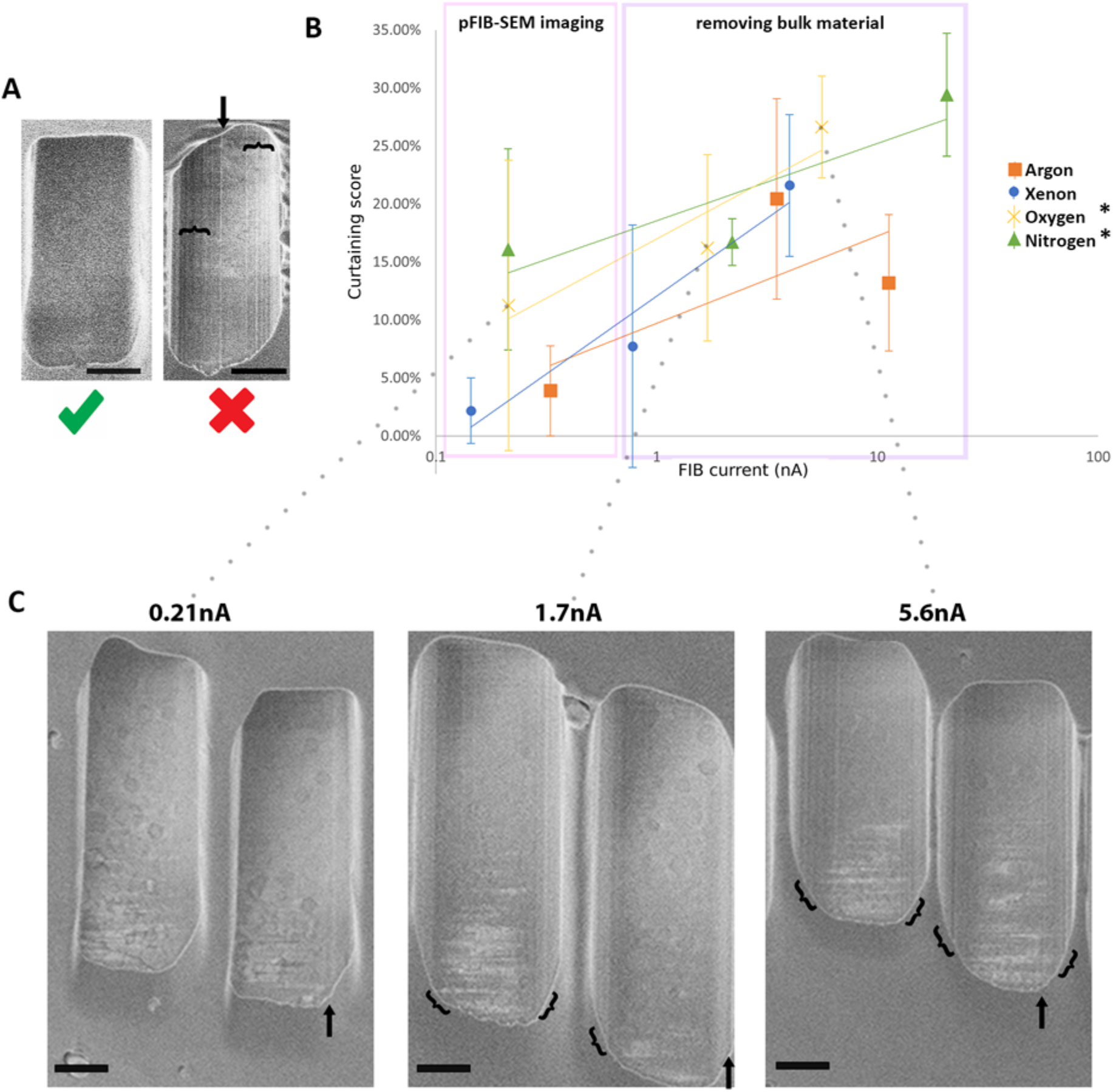
The curtaining score (higher equates to a greater curtaining propensity) for the different plasma sources at different currents. Plunge frozen *C.trachomatis-infected* HeLa cells were milled for each current and each gas. 15 windows of 2×2.5×2μm were milled at different measured currents at 30kV acceleration voltage for xenon, oxygen and nitrogen and 20kV for argon gas (see Figure 1 Supplementary Figure 2 and 3). The incidence angle was set to 18°. SEM images were acquired 90° to the FIB. (A) Representative images from these data, showing little or no curtaining (left, oxygen, 213 pA)) and extensive (right, nitrogen, 2.2nA) curtaining. Arrow and curly brackets indicate the position of a curtain or group of curtains. Scale bar: 1μm (B) Plot of curtaining score as a function of current (see materials and methods). Points represent the mean value associated with the standard error. (C) Representative images of windows generated with oxygen. Arrow and curly brackets indicate the position of a curtain or group of curtains. Scale bar: 1μm. **Supplementary figures: Figure 1 - supplementary figure 1; Figure 1 - supplementary figure 2; Figure 1 – supplementary figure 3; Figure 1 – supplementary figure 4.**

In addition, argon and xenon are monatomic gases which give rise to single plasma species (Ar^+^ and Xe^+^) whereas nitrogen and oxygen are diatomic gases producing multiple ion species (O^+^, O_2_^+^ and N^+^, N_2_^+^) which can lead to the formation of two images which partially overlap. For oxygen a double image is always formed irrespective of the imaging mode. With nitrogen, we observed the formation of double images when imaging in ‘immersion’ mode. (Figure1 - supplementary figure 4). This double image caused issues tracking the milled surface during serial pFIB/SEM, limiting the volume that could be obtained and leading to abortive runs. The double image can be compensated for via an extra lens in the FIB column, however, this compensation would have to be manually varied during the experiment.

We concluded that argon is the optimal gas to perform serial pFIB/SEM and we performed most of our data acquisition with. For completeness we also present data using nitrogen plasma, which has both double imaging and a high tendency for curtain with respect to the other gases.

### Initial imaging of cells and tissue

For our first experiments we decided to use the machine with the geometry as is, where the SEM image is formed at 52° relative to the FIB milling plane. We analysed human HeLa cells and intact tissue from the mouse (*M. musculus*) brain cortex. We used a milling step of 50 nm. This step depth was chosen based on a compromise between workflow and electron interaction depth. The interaction depth was informed by the work from Haase and colleagues (Guehrs et al., 2017) who estimated this to be 30 nm for entirely biological polymers and whilst imagining of silver beads inside cells reported an interaction depth of roughly 90 nm (Seiter et al., 2014). The focus was adjusted manually to the centre of the field of view (FOV).

The physical basis of image formation in SEM of flat surfaces (topology free) is based on the detection of secondary electrons and the contrast is formed due to the local difference of potential of different cellular constituents (Schertel et al., 2013). In our hands the best imaging parameters are between 1.1 and 1.2kV (low kV), with a current between 6.3 and 13pA (low current). A short dwell time (100ms) associated to line integration (50-100 time) are optimised settings on our system for optimal contrast and low charging artefact.

We recorded 500 slices on plunge frozen HeLa cell and clearly visualised various membranous compartments, such as nucleus, endoplasmic reticulum (ER), mitochondria, and mitochondrial cristae were all easily spotted (Figure 2A -B). We observed two centrosomes, recognisable with their respective pairs of centrioles characterised by their archetypal barrel/wheel shape associated to a pericentriolar matrix (Figure 4 C-E). The distance between the centre of the two centrosomes is approximately 1.2 μm consistent with the cell in the prophase (Figure 2 E) (Kaseda et al., 2012). The spacing of the centrioles are 543 nm (centre to centre) in one centrosome and 1561 nm in the other, a heterogeneity which is not uncommon for cells in division (Figure 2 E) (Vitiello et al., 2019). However, the spindle microtubules were not observed between the two centrosomes.

**Figure 2.**
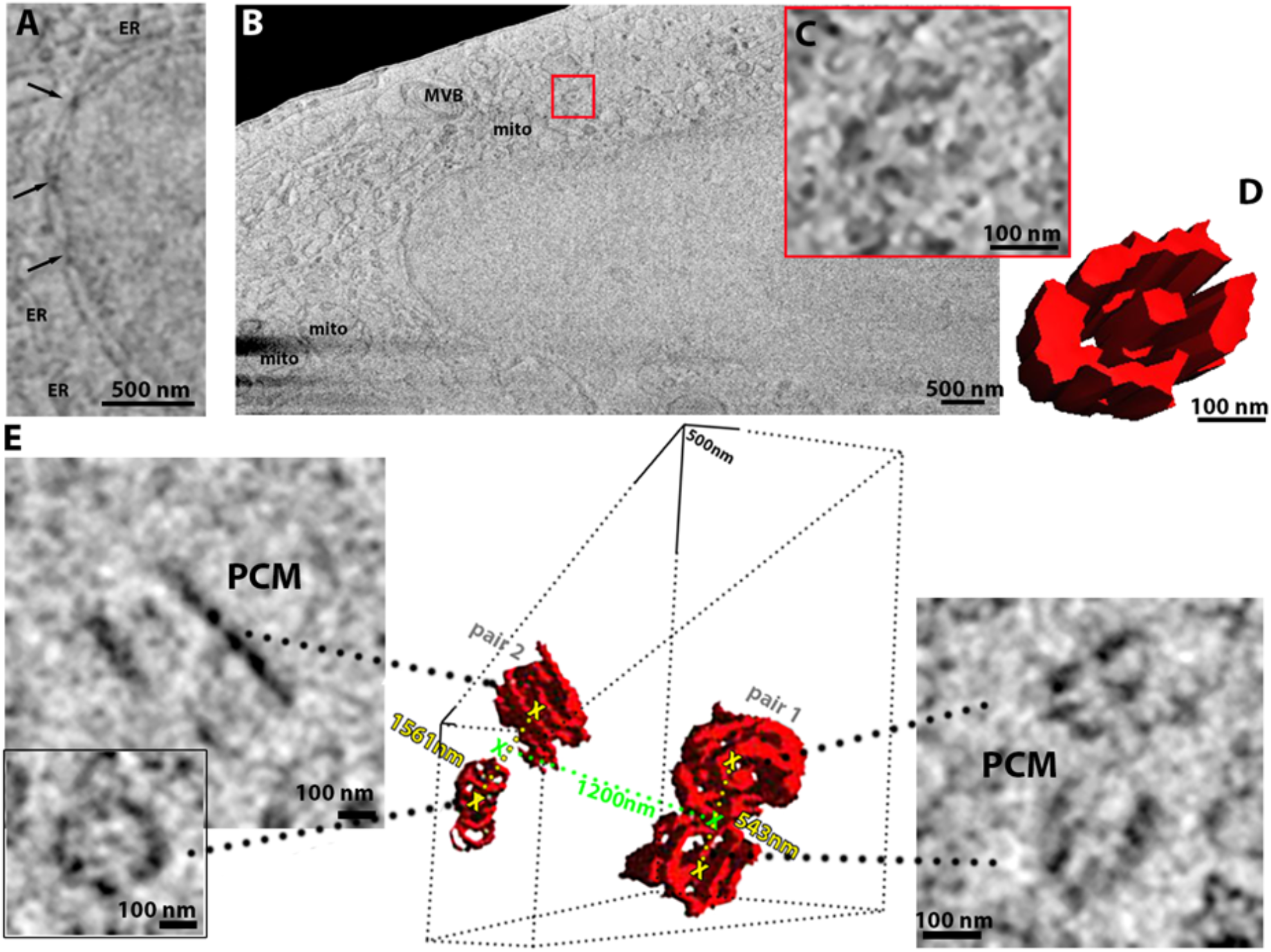
HeLa cells imaged using serial pFIB/SEM. (A-E) Serial pFIB/SEM volume acquired of a HeLa cell, using argon for milling and were imaged 52° to the SEM. (A) Zoomed in showing nuclear pore complexes (NPC, arrows) and endoplasmic reticulum (ER). (B) Overview where the nucleus, mitochondria (mito) and multivesicular body (MVB) are easily identifiable as well as a centriole (red box). (C) zoom of the centriole identified in (B) with its 3D rendering (D). (E) This HeLa cell presented two centrosomes with two pairs of centrioles and associated pericentriolar matrix (PCM). The distance in green is between the two centrosomes respective centres and the distance in yellow depicts the distance between the centrioles-in each pair. (D-G) Slices were filtered using a 2 pixels radius mean filter. For (E) we used a band pass filter. Full data are shown in Figure 2 - Supplementary Movies 1

**Figure 3.**
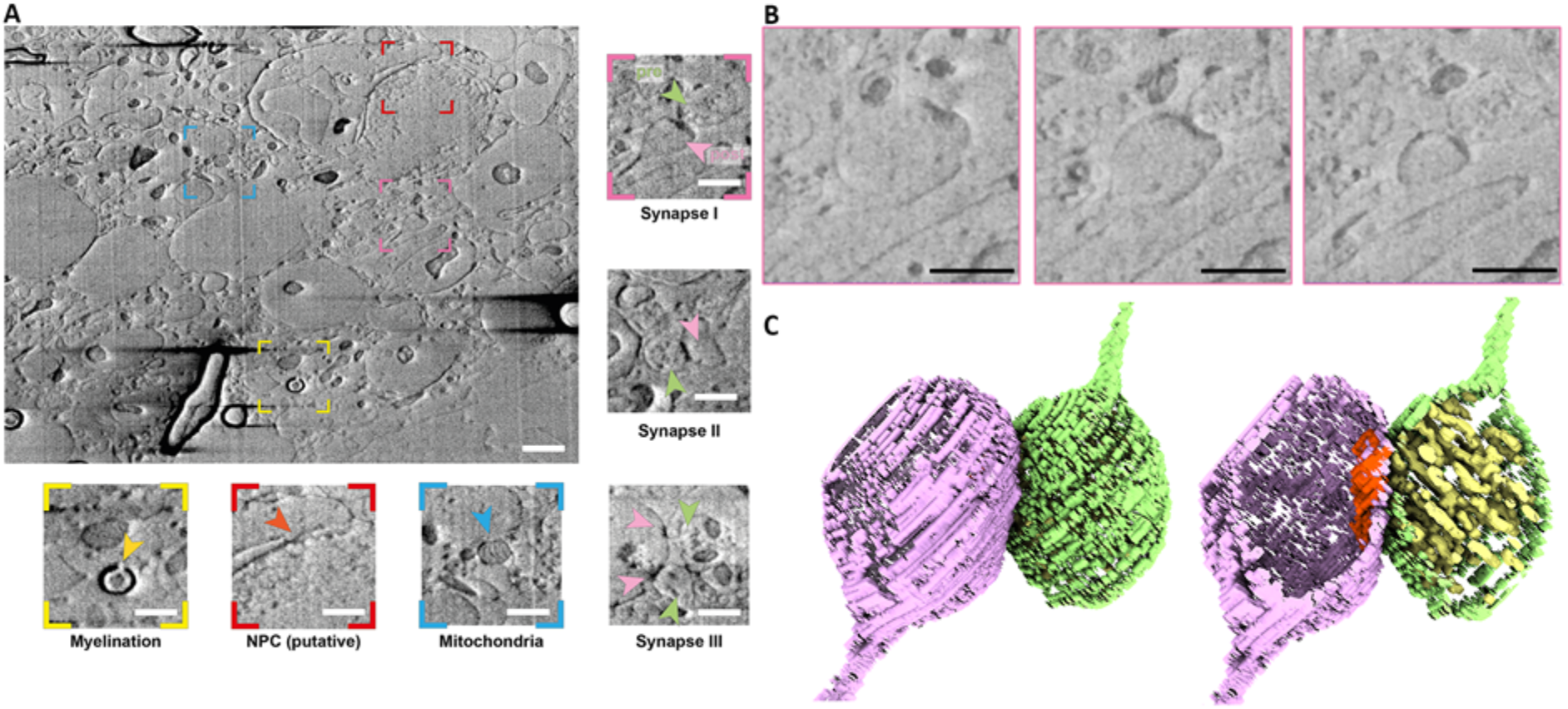
Non-fixed, HPF vibratome slice from a mouse brain milled using argon and imaged 52° to the SEM. (A) Representative slice of wide field of view of the serial pFIB/SEM volume. Scale bar: 1μm. Coloured insets show regions of interest within the field of view, including myelin sheaths (yellow), putative nuclear pore complexes (red) and mitochondria (blue). Non-coloured insets show synapse morphologies from different slices. Pre- and post-synaptic cells are shown with light pink and magenta arrows, respectively. Scale bar: 500 nm (B) Enlarged slices from region of (A) indicated in pink of a region containing a neuronal synapse, slices through Z shown at progressive positions, from left to right. Scale bar: 500 nm (C) 3D volume rendering of the synapse presented in (B) with the pre- (green) and post- (purple) synaptic membranes labelled. The post synaptic density (red) and pre-synaptic vesicles (yellow) are clearly identifiable. For presentation purposes the presented slices have been filtered using a 2 pixels radius mean filter. Full data are shown in Figure 3 - Supplementary Movie 1.

**Figure 4.**
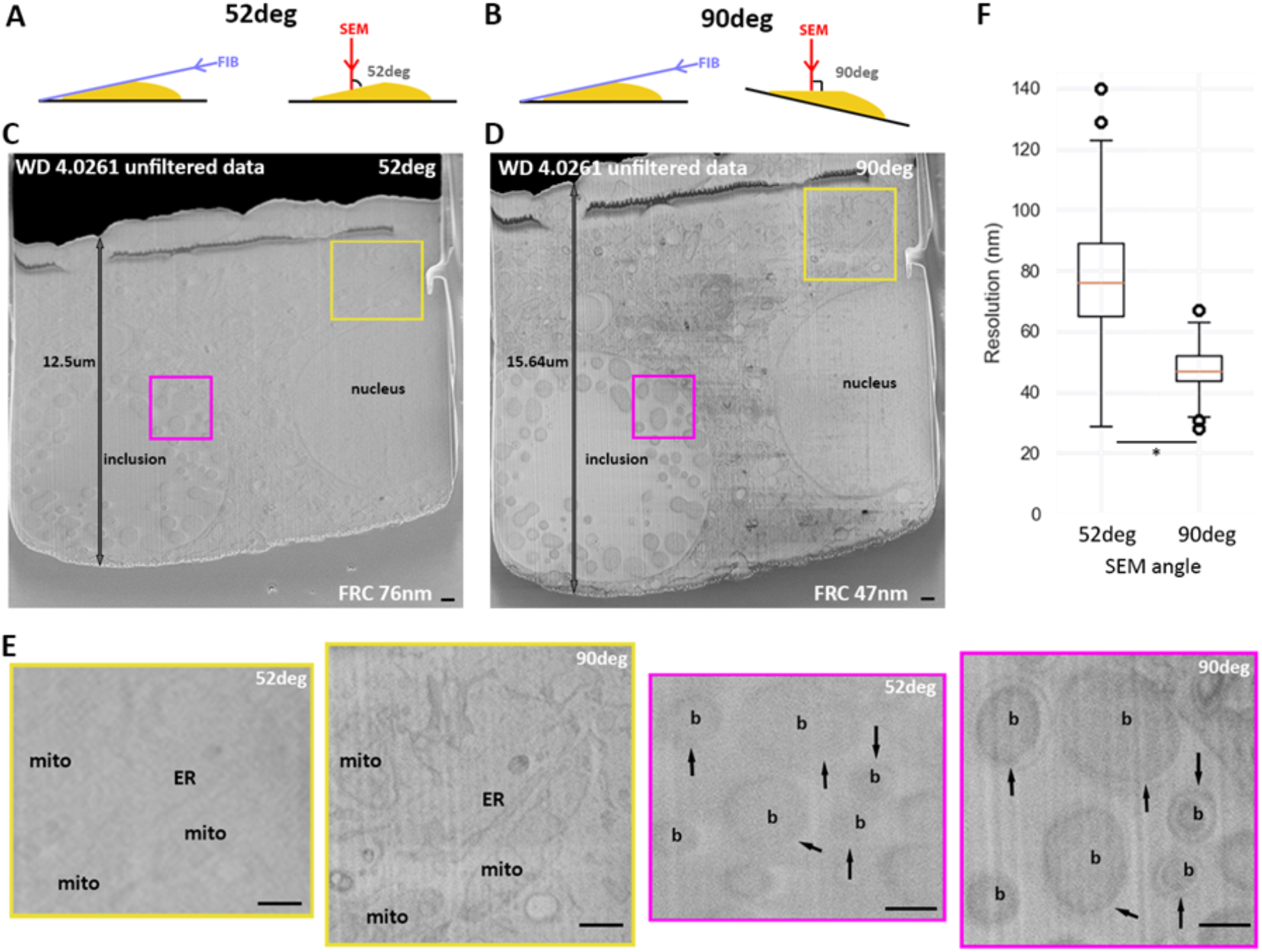
52° vs 90° stage tilt. **(A)** and **(B)** Schematic of SEM (red arrow) and FIB (purple arrow) angles during serial pFIB/SEM imaging at 52° and 90°, respectively. (A -E) HeLa cells infected with *C.trachomatis*. Images have been acquired with exactly the same parameters but the tilt angle. No histogram modifications have been made and no filters have been applied. (C) and (D) are the full field of view and (E) zoom in panel (colour boxes). Scale bars: 2μm (C and D) 500 nm (E). Arrows show the outer membrane of bacteria. WD: working distance, FRC: Fourier Ring Correlation, mito: mitochondria, ER: endoplasmic reticulum, b: bacteria. (F) presents the resolution obtained for the different patches on images acquired at 52°deg or 90°deg to the SEM using FRC to measure. **Supplementary Figures: Figure 4 – supplementary 1 to 5.**

To allow vitrification of tissue samples high pressure freezing (HPF) methods are needed as samples are too thick to vitrify via plunge freezing methods. HPF slices of mouse brain cortical tissue were prepared and imaged via serial pFIB/SEM (Figure 3). The brain tissue imaged volume was 844.4 μm^3^ (20.7 μm x 13.8 μm x 3 μm) and therefore permits observation of a range of cell types including neurons and oligodendrocytes. Cellular interactions were observed, including the envelopment of axons by oligodendrocytes (visible as black contrast in images due to the high myelin (lipid) content). These features appear as elongated (oblique plane milling) or ‘horseshoe’ (transverse plane milling) features. The diameter of these myelinated neurons ranged from ~0.53 to ~2.8 μm (longest cross section, mean 1.02 ± 0.73 μm) demonstrating that these can be highly variable. Within cells we clearly observed nuclei and mitochondria, including 87 cristae (Figure 3 A). Other membrane-bound compartments of varying size and diverse morphology were identified within the tissue (Figure 3 - supplementary movie 1). Synaptic junctions were visible, and we were able to discern the presence of pre-synaptic vesicles (typically 20-30 nm) within the pre-synaptic terminal. Within one terminal, ~ 184 vesicles were identified (Synapse I, Figure 3 A to C), whereas in a second, ~ 98 were identified (Synapse II, Figure 3 A). The resolution did not permit the internal volume of vesicles to be described. On the post-synaptic neuron, the well-known post-synaptic density could be visualised (Figure 3) (Valtschanoff and Weinberg, 2001). Both terminals led to extruded (thinning) membranes leading toward axons and dendrites, respectively. Interestingly, the morphologies of the synapses were very diverse, sometimes with multiple pre-synaptic terminals forming on one post-synaptic face (Figure 3A), highlighting the diversity prevalent in synaptic junctions in tissue. Such diversity is consistent with previous studies (Santuy et al., 2018).

### Optimising imaging quality

We suspected that we could further improve image quality by carrying out the SEM imaging 90° to milled sample surface. To achieve this, a compensatory stage tilt between milling and imaging steps is required (Figure 4 A and B). To evaluate the gain introduced by this tilt, we imaged HeLa cells infected with *Chlamydia trachomatis*. These cells can be readily plunge frozen and exhibit a wide range of sub-cellular features (bacteria, vacuoles, Golgi apparatus, nuclei, large protein assemblies, e.g., NPC) enabling visual analysis of contrast and information content. The samples are also large enough to enable a wide FOV. The cells were milled with Ar and subsequently imaged at the two different angles. Other parameters, such as stage positions, working distance and imaging parameters (voltage, current, line integration, exposure, detectors voltage offset and gain) were kept constant. The contrast improvement when SEM images are taken at 90° (Figure 4 C to E) is immediately obvious to the eye. Features within images acquired at 52° are less pronounced compared to imaging at 90° to the surface, with features such as the nucleus, cytoplasmic vesicles, and bacterial inclusions, while discernible in both angles, being much more prominent in images acquired at 90°. The outer membrane of the bacteria exemplifies a case where features are very low contrast when imaged at 52°, but which at 90° can be prominent seen in the images (Figure 4 E arrows). The dramatic shift in quality would be expected to have profound effects in image analysis and segmentation.

Imaging a tilted surface causes elongation of features along the y-axis in the field of view (Figure 4 A and B). We segmented the *C.trachomatis* cell envelope to quantify the effect of 52° tilted data on the sphericity (Figure 4 - supplementary figure 1 A and B). At 52° this elongation results in cells appearing as ellipsoids along the Y axis (Figure 4 -supplementary 1 C). At 90° these features are regular circles.

Although obvious visually, we sought to determine a more quantitative assessment of the improvement in image quality. This was achieved by implementing a Fourier Ring Correlation (FRC) approach based on the one-image FRC that is used in fluorescence microscopy (Koho et al., 2019) (Figure 4 - supplementary 2). In one-image FRC a single image is split into two by taking even and odd pixels in a checkerboard pattern and comparing the two. Patches of 256×256 pixels on the image were generated for assessment, producing maps of local resolution across the whole FOV (Figure 4 - supplementary figure 3).

In estimating the resolution improvement due to the difference in the angle, other imaging parameters were kept constant (Figure 4 C and D). SEM imaging at 52° produced a global image resolution on average over a whole FOV of 76 ± 19 nm, while using the same assessment with SEM imaging at 90° gave a resolution of 47 ± 7 nm over the whole FOV (Figure 4J). Manual imaging at 52° with optimization of focus improved the reported resolution (Figure 4 - supplementary figure 3) (53 ± 16 nm). Indeed, the bacterial outer membrane is now more visible at 52° than before. However, visual analysis of features including the outer membrane show that even with manual control, 90° imaging yields more detail and higher contrast.

Such stage movement can introduce image shifts which in turn can introduce complexity and errors in downstream analysis. We measured the XY and Z shift using 1 μm beads that were imaged at 52° (no movement) or at 90° following stage movement. The XY shift was calculated as the shift between features and has been measured to be 52 ± 38 nm without stage movement, and 96 ± 71 nm with stage movement. Thus, stage movement results in 2-fold increase in drift, however the absolute magnitude of the drift is small, representing under 1% of the FOV (Figure 4 – supplementary figure 4 A and B). This small increase in drift will have little (if any) effect on the alignment. For potential Z-shift, we used the sphericity (and thus change in diameter with depth) of the beads to determine discrepancies between the programmed milling step size and the actual step size (derived by measuring diameters). For a targeted milling step of 50 nm, the observed steps were 48.1 ± 1.4nm without stage movement and 48.9 ± 4 nm with stage movement. Z-drift is minimal in both cases and not increased by the tilting regime (Figure 4 - supplementary figure 4 C and D).

The depth of field is of interest as it informs how much volume can be milled without the need for autofocus correction methods whilst retaining image quality. At both 1 and 2kV imaging energy, this was found to be ~20 μm (Figure 4 - supplementary figure 5). Using a slice thickness of 50 nm, a focus correction will be needed after around 400 slices. Fully automating focus corrections would be desirable for thicker samples. Alternatively, a focus step change based on the milling steps could be implemented (Auto Slice and View, ASV, from ThermoFisher).

### Optimal cryo serial pFIB/SEM

We applied the 90° imaging approach to a range of samples to evaluate its potential to generate new biological insights.

### R. rubrum

*R. rubrum* are purple bacteria which present a more complex cytoplasmic organisation due to the photosynthetic capabilities of this organism. Chromatophores are organelles supporting photosynthesis and their formation have not been observed in *R. rubrum*. Plunge frozen purple photosynthetic *R. rubrum* were imaged in a frozen-hydrated state, absent of stain or fixative (Figure 5). Minimal sub-cellular artefacts were observed, and biological features including lipid droplets and inner and outer membranes, postulated to be 30-80 nm apart (Tucker et al., 2010) were clearly distinguished within the pFIB/SEM image stack (Figure 5 A-B). Collection of the dataset was accomplished in 2.5 hrs in an automated manner. We observed both developing and mature chromatophores; the roughly spherical vesicles which contain the photosynthetic apparatus and are ~30-60 nm in diameter when mature. For the first time, nascent chromatophores budding from the inner membrane were unambiguously discerned (Figure 5 A). For the 8 cells which we segmented, 2772 mature and 84 nascent chromatophores were observed. The segmented data comprises ~33% of the total number of cells, hence an estimated ~8000 chromatophores would be available for analysis in the full volume. We noted that the nascent chromatophores were unevenly distributed; in one of the segmented cells, ~12% of the total chromatophores were nascent whereas the majority of cells imaged had very few or none. The growth conditions and sample preparation used were not favouring the replacement of photosynthetic machinery. The identification of this atypical and rare cell was possible as we imaged several bacteria volumetrically at excellent mesoscale resolution. The ability of serial pFIB/SEM ability to image large volumes quickly allows rare events such as chromatophore budding to be observed.

**Figure 5:**
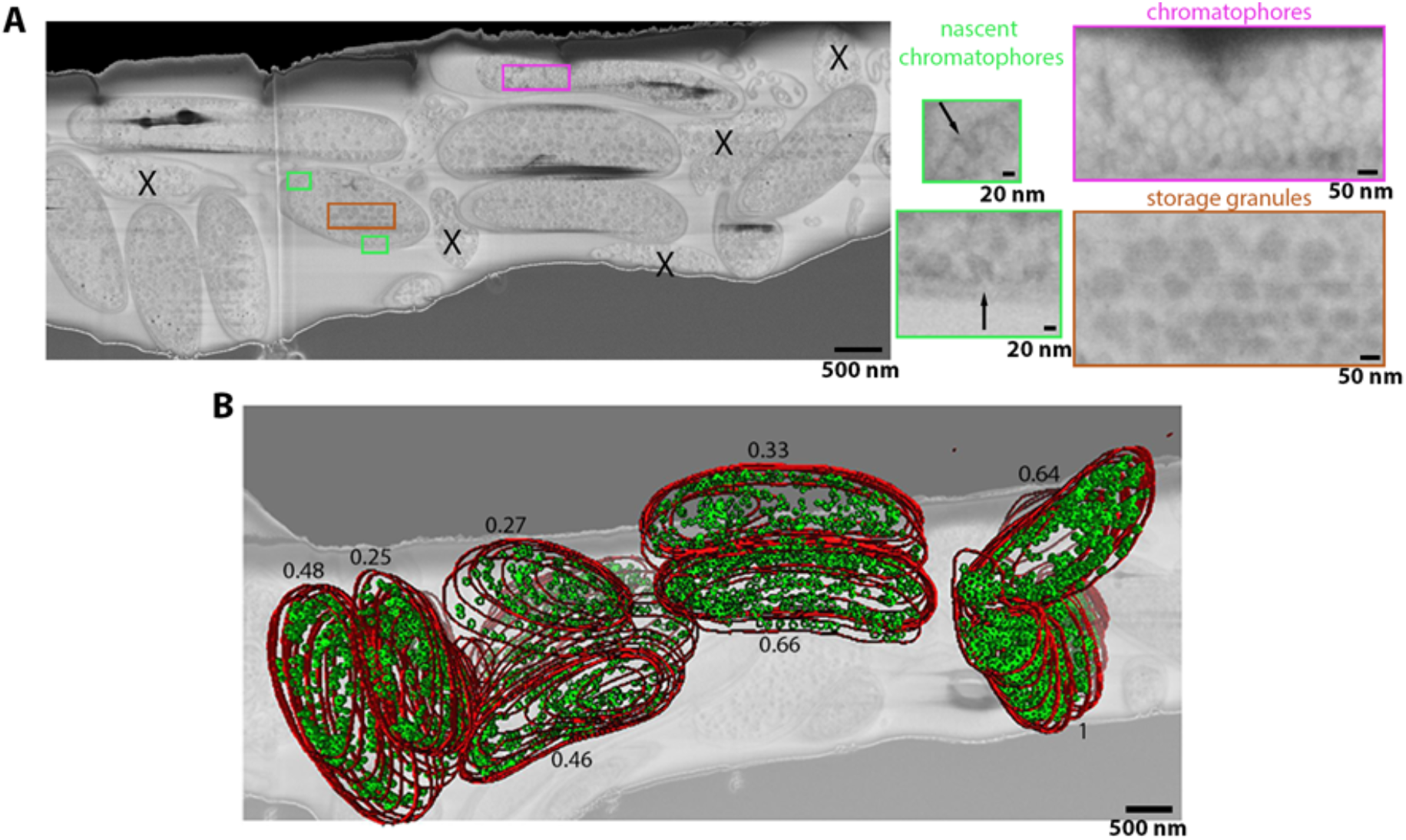
*R. rubrum* image stacks acquired using argon serial pFIB/SEM normal to the SEM column. (A) Full field of view of slice of serial pFIB/SEM volume, showing vitrified *R.rubrum*. Features within the bacteria are shown enlarged (right), with highlighted areas from showing nascent chromatophores (green), mature chromatophores (pink) and storage granules (brown). X indicate dead/dying bacteria or debris. (B) Slice of the volume superimposed with the volume rendering after segmentation. In red is the outer membrane and green the mature chromatophores. The number associated with each segmented bacteria is the ratio of the number of chromatophores per sum of the surfaces occupied by the bacteria at each slice. The slice was filtered using a 2 pixel radius gaussian filter. Full data are shown in Figure 5 - Supplementary Movie 1.

### Vero cells

Vitrified Vero cells (plunge freezing) were milled using nitrogen. Nitrogen was chosen to investigate whether the worst performing gas (in terms of curtaining) was still able to provide useful insights. Imaging with nitrogen severely limited the software’s ability to track the milling area, this is reflected in the number of Z slices acquired compared to HeLa cells (46 slices vs 500 for HeLa). This volume was also acquired after 7 previous attempts to automate serial FIB/SEM acquisition. However, we were able to clearly see sub-cellular structure. In addition to membranous organelles, we could identify large protein complexes within the cellular volumes. Nuclear pore complexes (NPCs) could be observed (Figure 6 A) within cells. This enabled us to visualise their organisation within the cell, with 138 NPCs being visible within the data, for example. The expected mitochondrial network (Shen et al., 2022) was also characterised (Fig 6 A, B and D) and both isolated mitochondria (mitochondria 16 and 17), small mitochondrial networks (12 and 14), and large networks (2 and 7) could be identified. Additionally, we could calculate the volume taken up by mitochondria, ER and lipid droplets within the volume we interrogated (Figure 6 A, B and E). We could also quantify the number and size of contacts of the ER with lipid droplets (LD) or with mitochondria (Figure 4 A, B and E). In Vero cells, we identified 26 LD-to-ER contacts and 48 mitochondria-to-ER membrane contacts sites (Giacomello and Pellegrini, 2016, Jacquemyn et al., 2017)). Membrane contact sites (MCS) vary in length (membrane to membrane contact) and distance between the two organelles. In our analysis, we kept a constant distance between organelles (25 nm which is the largest space between membranes). Therefore, the analysis is independent of the intermembrane space variation and includes all MCS. Both LD-ER and mitochondria-ER MCS have a similar volume (1 to 5 x 10^-3^ μm^3^); (Fig 6 A) and gaussian distribution. Consequently, ER contact sites measures vary depending on how long and how spread are those contacts. Interestingly, the distribution is the same, suggesting a limit in the extent of which the ER can track and contact an organelle.

**Figure 6.**
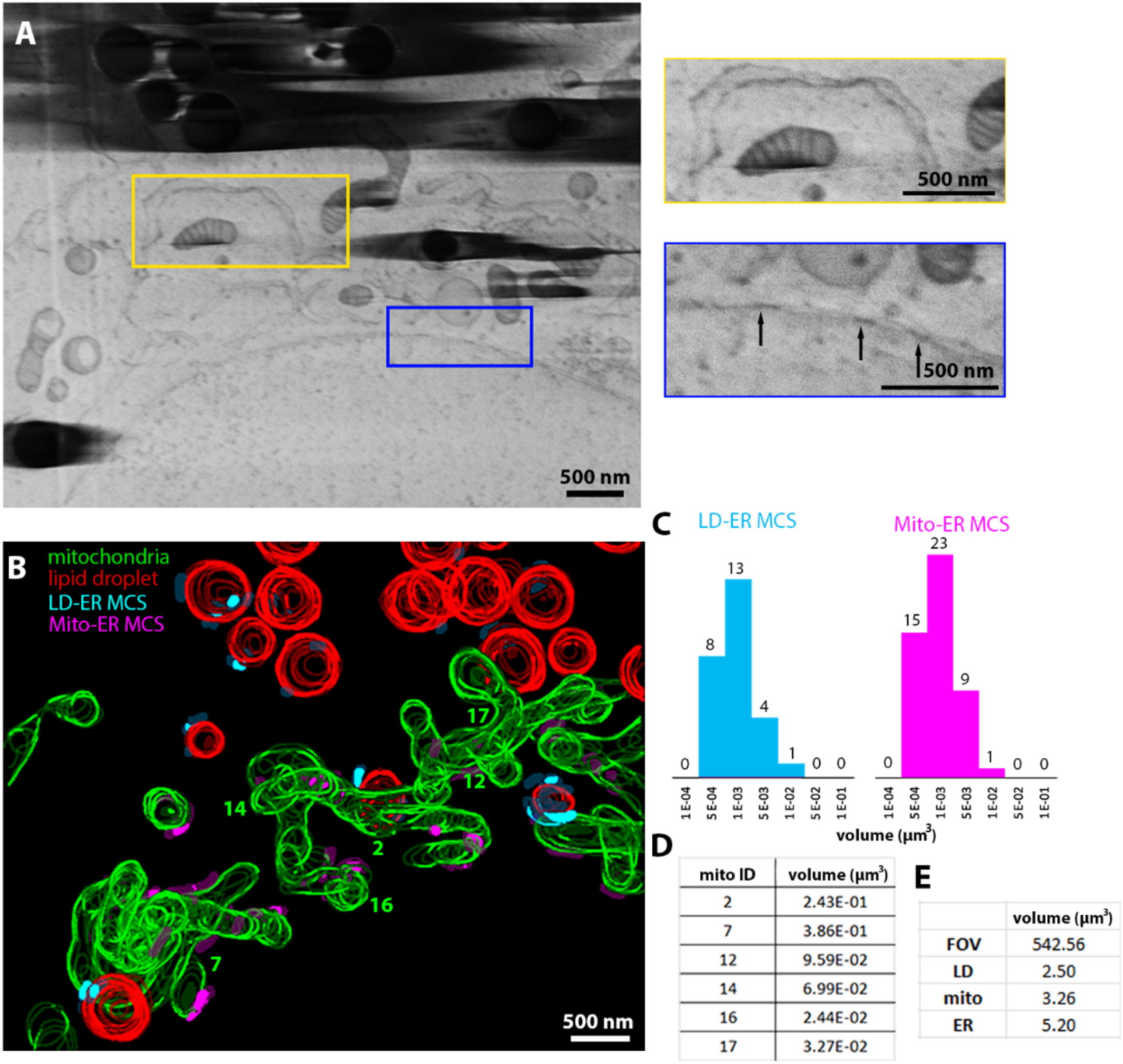
Vero cells imaged using serial pFIB/SEM. (A-E) Serial pFIB/SEM volume acquired of a Vero cell, using nitrogen for milling and were imaged 90° to the SEM. (A) Slice of volume of a Vero cell imaged 90°. Insets show enlarged regions of endoplasmic reticulum (ER) and mitochondria with visible cristae (yellow), and enlarged region populated with nuclear pore complexes (blue) (arrows). N.B. presence of curtains towards the left of the yellow box in the large FOV image (left) due to contamination at the surface of the cell. (B) 3-D segmentation of subcellular features within the volume from Vero cells presented in (A). Red: lipid droplets (LDs), green: mitochondria, cyan: LD to ER membrane contact site (MCS) (max 25nm space between the membranes), purple: mitochondria to ER MSCs. (C) Number of different ER MSCs vs volume size for contact with lipid droplets (blue) and mitochondria (pink) (μm^3^). (D) Volume (μm^3^) of complete mitochondria within the volume of the Vero cell is shown. (E) volume (μm^3^) of the field of view (FOV) and different organelles. Full data are shown in Figure 6 - supplementary movie 1 and full segmentation including ER are available on Figure 6 - supplementary movie 2.

### High pressure frozen heart tissue

To stop the heart beating while maintaining tissue integrity, we first fixed with 4 % PFA prior to sectioning and vitrified a mouse heart tissue. Cryogenic serial pFIB/SEM was then subsequently performed without any other treatment and vitrified in similar manner to the brain tissue. Sarcomere-containing regions (myofibrils) could be visualised sandwiched between columns of mitochondria, running along and at oblique angles with respect to the milling direction (Figure 7 A). The Z-disk, I-band and A-bands of the sarcomeres were visible, along with a putative M-line (Figure 7 A and B). The average sarcomere length in these samples was 1.68 μm (SD=0.03, N=34,). As cardiomyocytes are syncytial, they have multiple nuclei. One nucleus was seen with associated nuclear pore complexes and putative hetero-/euchromatin regions. Blood vessels, in proximity to the cell, were identified by the characteristic layer of endothelial cells and red blood cells (bi-lobal shape) (Figure 7 B). In the heart volume examined (2698.4 μm^3^), 997 mitochondria were visualised. This gives a relative concentration of mitochondria of 0.36 mitochondria/μm^3^ for heart tissue, (c.f. 0.10 for mouse brain), which agrees with previous observations of mitochondrial content in mouse heart tissues (Else & Hulbert, 1985).

**Figure 7.**
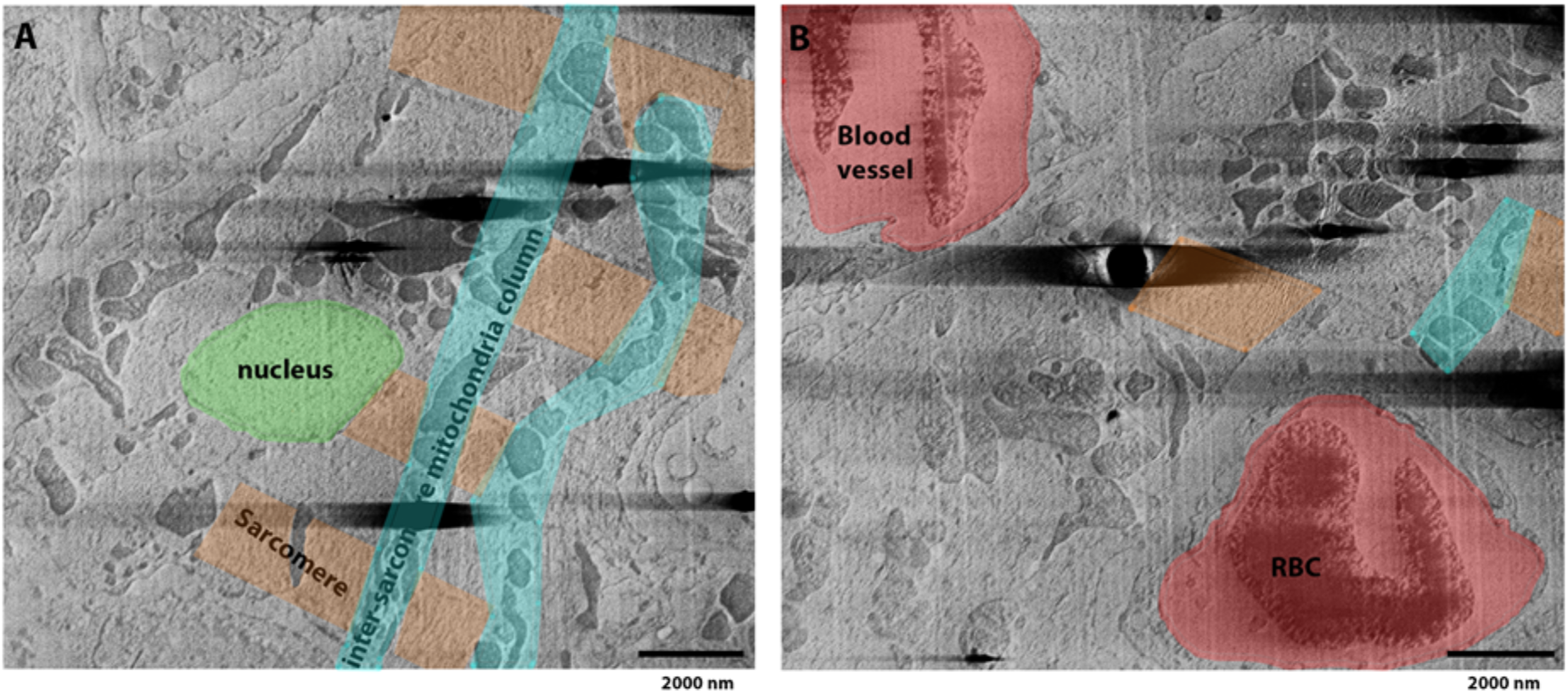
Fixed, HPF vibratome slice from a mouse heart image milled using argon and imaged 90° to the SEM. (A) Slice from serial pFIB/SEM volume showing sub-cellular structures consistent with those recognised from a cardiac myofibril (Begay et al., 2018), with a nucleus (green), sarcomeric elements (including Z-disk, I-band and A-band) (orange) and mitochondria organised between axially organised sarcomeres in columns (cyan). (B) Slice above the image shown in (A), where amongst the cardiac tissue, blood vessels and red blood cells (RBC) can be identified (red). Figure 7 -supplementary movie 1 presents the whole data.

### Resolution estimation of the datasets

The FRC approach was used to assess the resolution of our datasets. Interestingly, the brain and HeLa cells which have been acquired using the same software, angle and pixel size present different resolutions reflecting the importance of other parameters such as focusing, stigmation as well as sample to sample variability. *R. rubrum* and Vero cells also have been imaged using similar pixel size, identical software and imaging angle and present a difference in resolution (2-fold in favour of nitrogen). The FRC measurement of images at the beginning, middle and end of each of the biological sample datasets was calculated to allow assessment as to how resolution may vary over a given serial pFIB/SEM acquisition series (Figure 8 and Figure 8 - supplementary figure 1). We determined that in each acquisition series there is no indication of resolution degradation over the course of data collection, even in the case of longer data acquisition runs (HeLa cells - 500 slices). In this case the software used was ASV which integrated estimation-based focus change for each step.

**Figure 8.**
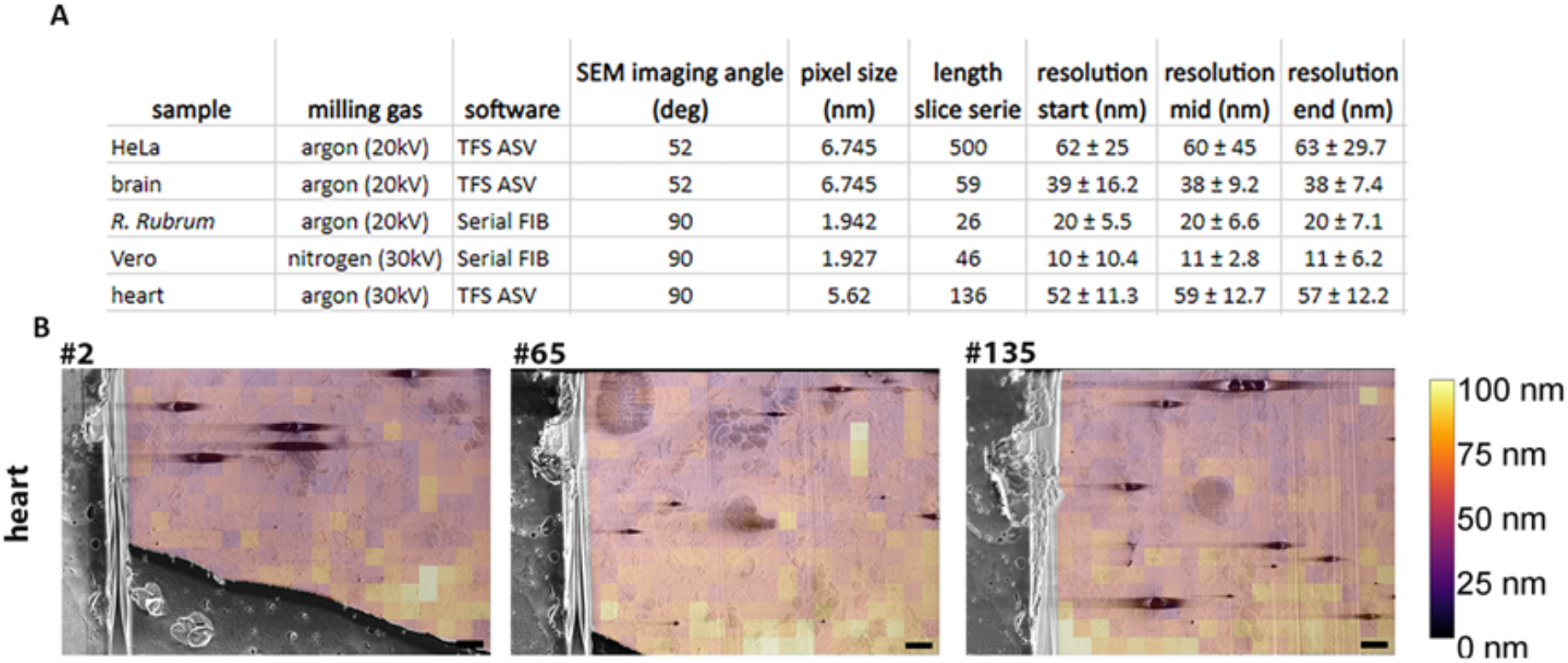
Resolution measurements for biological samples -. (A) Table presents the acquisition parameters and the associated calculated resolution (FRC calculation) for the dataset presented in Figures 3 to 6. The length of the series represents the number of slices. All datasets were acquired using a 50 nm step. Slices from serial pFIB/SEM volumes FRC resolution measurements were taken using three slices (start, mid, end) from each dataset. In all cases, the GIS layer was masked out of these measurements. Software used are either SerialFIB (Klumpe et al., 2021) or ThermoFisher Scientific Auto Slice and View (TFS ASV) (B) example of the FRC analysis from the heart dataset. Slice number 2, 65 and 135 are the one used for start, mid and end. Scale bars: 2μm. Figure 8 -supplementary figure 1: FRC overlays for HeLa, Brain and *R. rubrum*.

### Mitigating charging artefacts

Charging occurs when the energetic electron beam interacts with highly insulating substances in the sample, which is consistent with our data where charging was typically found in regions associated with a greater lipid content (lipidic membranes, lipid vesicles, myelin) (Figures 2–7). The presence of charging manifests as intense dark spots with asymmetric streaks in the direction of scanning.

We implemented an algorithm that enabled computationally removal of the charging associated with an assessment of the underlying, non-artifactual features within the image within the data. *S. cerevisae* data imaged at 90° to the surface was used due to the presence of extensive charging of lipid droplets. While enabling us to qualitatively characterise the effect of charging on image content, such approaches may also facilitate further segmentation and image analysis. The process required segmentation of the charged centres (not including the asymmetric streaking) using a pre-trained U-Net neural network (Ronneberger et al., 2015) with slight manual clean-up (Figure 9 A). Comparing the automated segmentation of charge centres with the automated with manual clean-up yielded a 0.93 Dice coefficient score (Dice, 1945; Sørensen, 1948). This high Dice coefficient score indicates little manual clean-up was required and full automation of this step may be possible in future.

**Figure 9.**
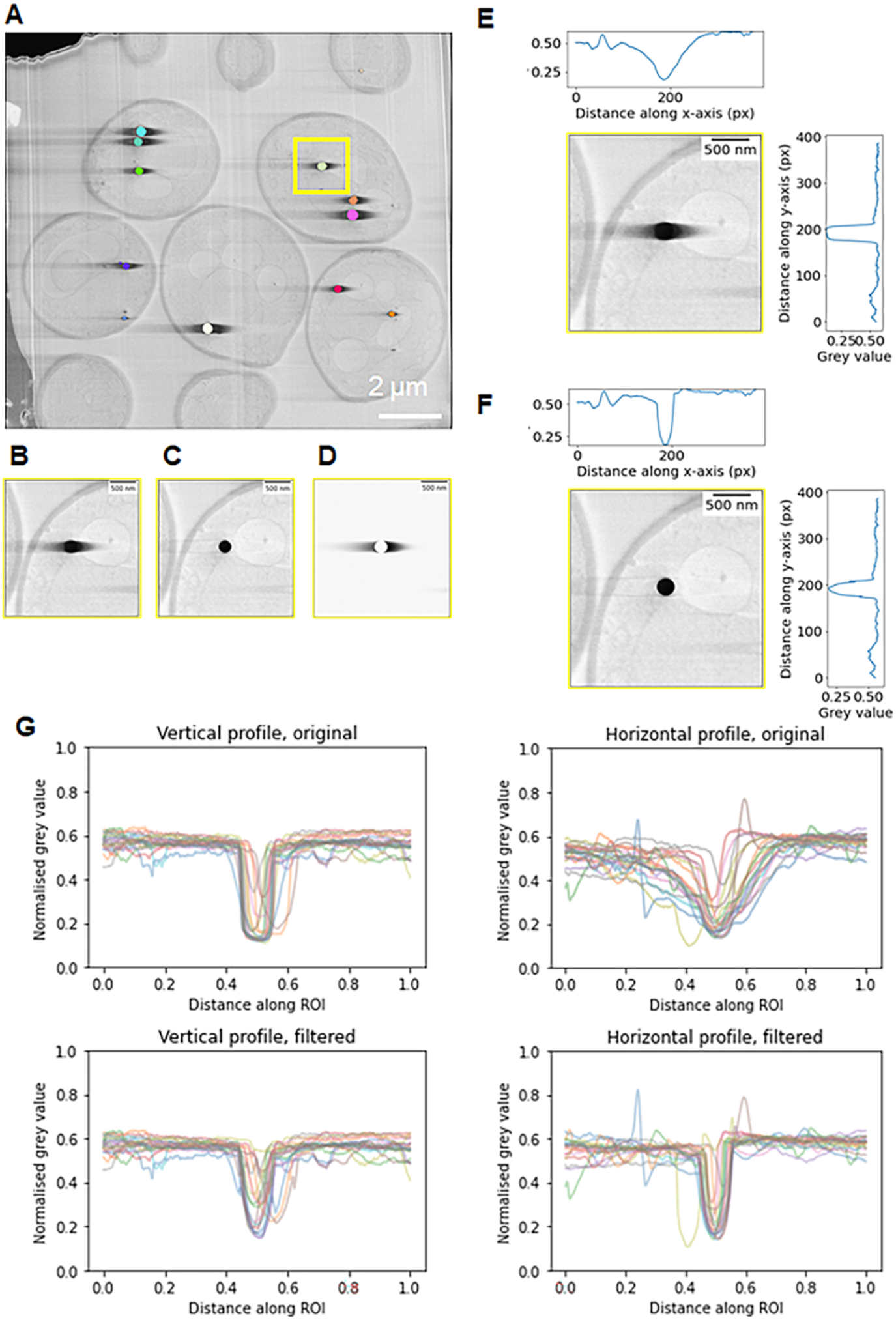
Post-processing approach to mitigate charging artefacts. Charging artefacts around lipid droplets in *S. cerevisiae* were removed and the effects measured. Cells were imaged at 90° to the SEM and argon was used for milling. (A) Instance segmentation of lipid droplets found in the dataset. (B) and (C) show the lipid droplet highlighted in the yellow box in (A) before and after charging artefact suppression. (D) shows the asymmetrical charging artefacts in isolation, given by (C) subtracted from (B). Because the artefact is asymmetrical, different functions were fitted on the left and right of the charging artefacts to locally mitigate this effect. (E) and (F) are individual example vertical and horizontal grey value line profiles through the lipid droplet in (B) and (C) respectively. (G) aggregated vertical and horizontal line profiles for 20 lipid droplets from (A) show that filtering to remove charging artefacts restores the sharp dip in grey values for the lipid droplets in the horizontal line profile, and more closely matches the grey values on either side of the charging centre, as can be seen in the vertical line profiles.

After charge centres have been segmented, a localised, row-by-row filter was applied which gathered the data near the annotated regions and compares it with the average of preceding rows. A smoothly varying continuous function is fitted and subtracted from the data line (see Materials and Methods). As a result, biological features adjacent to the charging artefact are partially restored (Figure 9 B and C). However, for regions immediately adjacent to the charged region itself, there is no information to recover. After application of this filter, the overall image resolution remains unchanged.

To quantify the effects of the charge mitigation, averaged line profiles were used in both the horizontal and vertical directions, centred on each charging centre before and after charge mitigation (Figure 9 E and F). For this analysis we used isolated complete charging centres representing. A line profile representing 9x the diameter of the charging centre was used to fully capture all effects due to charging (Figure 9 G). Prior to charge mitigation, the average contrast of the biological material adjacent to charging centres is much diminished and the dip in the line profiles due to this is variable and broad dependent on the size and severity of the artefact. This contrasts with the vertical line profiles where a clear, sharp dip due to the lipid droplet is consistently seen with grey values indicative of biological features on either side. Upon mitigation of the charging artefacts, a similar profile can be recovered, though with some asymmetry to the grey values still present on either side of the charge centre.

## Discussion

Imaging of biological materials using SEM is well known but challenging since biological material is primarily composed of light atoms (C, N, O) in a water background. This is traditionally overcome by some form of fixation and heavy metal staining which generates contrast (Knott et al., 2008; Watson, 1958). This is a very powerful approach and can be repeated using some form of ablation (ion beam, knife) to expose fresh surface in a serial process. Thus, volumes can be imaged in a series of steps. However, the process of fixation and staining by their nature alter the underlying biological structures (Thompson et al., 2016). Furthermore, these treatments are incompatible with further downstream high-resolution imaging by high resolution cryo-ET.

Building on work which applied SEM on vitrified hydrated samples (Schertel et al., 2013) we have evaluated different plasma gases for their ability to remove bulk biological materials while forming a smooth surface with minimal curtaining artefacts. Our work demonstrates that all four gases we used (O, N, Ar and Xe) (Figure 1) can under some conditions produce smooth surfaces that are amenable to SEM imaging. The precise behaviour of the different plasma species differs depending on the ion current used (Figure 1) and for Nitrogen on the acceleration voltage used (figure 1 - supplementary figure 2). Our results show that there is a difference in the curtaining propensity depending on the used gases and ion current, with N and O showing large curtaining propensity whilst Xe and Ar showed reduced propensity. We speculate that monatomic gases are less reactive at the surface of biological material, and less prone to curtaining.

We have shown that pFIB milling is effective for both plunge frozen cells and HPF mouse tissue. We observed a higher tendency to produce curtain artefacts in brain and heart tissue, something we attribute to the greater initial unevenness of the material due to the presence of lumen, facias, and supportive layers which mill differently. Here we used gaseous organic platinum to coat the sample. Since both composition and application method of the protective layer alter curtaining propensity (Leer et al., 2009), there may be gains from further exploration of different coating approaches.

The curtaining score that we have used is non-subjective but can miss some curtains likely leading to a consistent under assessment of the curtaining score. There is scope for this measure to be improved and, in future a version of the measurement that is applied on-the-fly could be used to automatically adjust milling parameters to prevent (and eliminate) curtaining during data acquisition.

The use of diatomic gases (O, N) as plasma source led to double images which can be (to an extent) corrected for but required more manual operation (Figure 6) and resulted in a higher failure rate for automation (under 20% success rate against more than 90% for argon). The use of oxygen plasma on resin embedded and stained samples has been reported (Gorelick et al., 2018) however the success rate was not reported. Since our goal was to develop automated procedures, we did not pursue these gases further. We observed that the propensity of Ar to introduce curtaining was similar to Xe at low current; thus, both seem strong candidates for exploitation in this workflow. At high currents Ar performed better, as demonstrated by its reduced curtaining score and lack of curtains when inspected visually. Consequently, we focussed Ar to demonstrate the potential of our approach.

The use of pFIB to fabricate lamella has been demonstrated with plunge frozen vitrified materials as part of a high-resolution high throughput cryo-ET workflow (Berger et al., 2022). The cryo-ET workflow study indicated that milling with Ar plasma (30kV, 60 pA) beam adversely affected the resolution of sub-tomographically averaged ribosomes to a depth of around 30 nm. This effect was present despite the “smooth” surface of the lamella and was attributed to damage introduced by Ar colliding with and penetrating the sample. The dependency of obtainable resolution on the ion source used has been shown in electron diffraction (Martynowycz et al. 2022). For Serial pFIB/SEM, the wattage used for argon 2-fold higher than that used for the final step of lamella polishing (200 pA at 20 kV vs 50 pA at 30 kV). Intuitively, we would expect damage (not just curtaining) to increase with increasing current, although this has not been formally proven. Interestingly when we compared cryogenic serial pFIB/SEM (low current) images of Vero (milled with nitrogen plasma) and HeLa (Ar plasma) cells, we measured a difference of 2-fold in resolution in favour of nitrogen. This hints that the non-curtaining damage may be dependent on the plasma not simply on its current and possibly the extent of this damage may not simply correlate with curtaining. Further investigation will be required. SEM imaging will damage the sample, the highest dose we employed data acquisition with the small pixel size was under 10^-2^ e^-^/Å, lower than the one used during cryo-ET imaging (Berger et al. 2022). Though damage increases at the acceleration voltage of SEM due to the increased scattering cross-section, the damage would be expected to be confined to the interaction layer, a depth less than 50 nm. Therefore, we expect SEM induced damage would be present in the same surface layer as the FIB damage and would not exceed the FIB damage layers. However, further experimentation may disclose unexpected effects.

Using low current, we were able to mill in a highly controlled manner, ablating 50 nm reliably and with high accuracy. However, the ideal depth of the milling remains unclear. SEM images are projections which lack depth information. Thus, in theory the ideal milling step will exactly match the depth of penetration (electron interaction) of the SEM; estimated to be between 20 and 30 nm (Kanaya & Okayama, 1972). Moreover, the depth of penetration is a function of the elements involved and will vary by sample. 50 nm was chosen as a compromise. Smaller steps require more time and potentially introduce cumulative electron damage. We did not notice obvious vertical discontinuities due to excessive electron dose (SEM) and features were tracked across the FOV in serial images. The rate limiting step in the automated workflow was the SEM imagining (6 - 7 minutes), with 30 s on average taken for milling of a 20 x 4 μm rectangle using Ar. Thus, using a plasma which mills faster at low power currently would have no significant improvement in the overall time taken for cryogenic serial pFIB/SEM. However, increasing the speed of SEM would be extremely beneficial. Scanning multiple lines concomitantly and/or software solutions (de Haan et al., 2019) would enable the analysis of larger volumes and would improve the integration of serial pFIB/SEM into other pipelines.

The quality of the resulting cryogenic serial pFIB/SEM images were excellent even with sub-optimal SEM imaging geometry. Using an algorithm derived from fluorescent imaging we calculated the local resolution across the image. This varies between samples and data acquisition parameters most importantly, and not surprisingly, the pixel size is critical. The patch size was maintained at 256×256, a smaller patch and more local analysis may lead to more “optimistic” resolution estimates. Our observation of protein features is consistent with the calculated resolution of < 50 nm.

By imaging 500 slices of 50 nm depth from HeLa cells, we were able to identify nuclear pore complexes and centrioles, both macromolecular complexes. We found two centrosomes indicating a cell in prophase but did not observe the spindle microtubules. In Vero cells, we delineated the mitochondrial network in exquisite detail and were even able to quantify the ER to organelle contacts. In mouse brain tissue, we visualised complete synapses including presynaptic vesicles (approx. 30-50 nm), identified myelin sheaths and cell cell interactions. Previous studies (Schertel et al., 2013) using SEM on high pressure frozen cryogenic samples have demonstrated the applicability of SEM to visualise sub-cellular structures, including the perinuclear space. Our data can directly identify large macromolecular complexes even within tissues, representing a significant improvement in the power of the approach.

In all samples, we did observe sample charging. This was most severe on heavily lipidated regions. We implemented a software mitigation approach that reduces the effect of charging on the surrounding data, though this was unable to recover information that was otherwise lost due to charging. Preventing the building up of such artefacts in the first instance would be preferable.

We improved the quality of the imaging by introducing extra steps such that SEM imaging occurred normal to the plane, which should be optimal. A systematic comparison of *Chlamydia* infected cells showed clear improvements in the definition of the ultrastructure. This was most clearly seen in the sharpness of the bacterial inner and outer membranes with the infected cells (Figure 4). The difference in quality between the imaging regimes could be reduced but not eliminated by manual optimisation of the working distance (Figure 4 - supplementary figure 3). The new imaging regime does require additional movements of the stage which for a highly repetitive process can lengthen the time taken and introduce compounding errors in stage position. However, the additional time was less than 1.5 % and we demonstrated that these additional movements have a negligible effect on the stage. Using this tilting approach, we examined photosynthetic bacteria, *R. Rubrum*. We were able to segment both mature and separately developing chromatophores. We also imaged unstained (but lightly fixed) mouse heart. We were able to measure the Z-disc to Z-disc distance 1.68 μm, which could reflect the heart being in diastole, but more likely demonstrates that the sarcomeres had undergone contraction post-mortem (where typical length in vivo is ~1.7 μm to 2.3 μm (de Tombe & ter Keurs, 2016). The images also allowed accurate assessment of the mitochondrial density in this tissue and quantitative comparison with the brain tissue. The ability to visualise protein assemblies and ultrastructure in vitrified unstained tissue with laboratory-based equipment has potential applications in pathology, where early identification of disease mechanisms remains an unmet challenge.

Cryogenic serial pFIB/SEM is a useful addition to the structural biology toolbox. We have shown that it can accurately determine biological ultrastructural arrangements from simple bacteria to complex tissue. The approach can generate a three-dimensional map of biological materials. The approach has several key advantages: it is highly controllable, it is relatively rapid, can be done in a laboratory and it is compatible with subsequent imaging modalities.

## Material and Methods

### Sample preparation

#### Mammalian cells

HeLa cells (CCL-2, ATCC, Manassas, VA, USA) were grown in high glucose DMEM with non-essential amino acids supplemented with 10% (v/v) foetal bovine serum (Gibco, ThermoFisher Scientific, Waltham, MA), 1% (v/v) Penicillin/streptomycin (Gibco) 1% (v/v) glutamine (200mM stock, Gibco). Vero cells (CCL-81, ATCC) were grown in DMEM (Gibco) with 1% FBS, 1% P/S and 25mM HEPES (Gibco). Cells were maintained at 37°C and 5% CO2 at >80% humidity. Cells were then seeded onto gold grids (either ‘UltrAuFoil’ 200 Au mesh, R2/2 gold film, or Quantifoil 300 Au mesh, R2/2 carbon film (Quantifoil Micro Tools, Großlöbichau, Germany) and grown to 50 % confluency before plunge freezing. Grids were then plunge-frozen in nitrogen-cooled liquid ethane using a Vitrobot (Thermo Fisher Scientific). Grids were then clipped into Autogrids (Thermo Fisher Scientific) and stored in liquid nitrogen.

#### Chlamydia trachomatis

The intracellular strict gram-negative bacteria *Chlamydia trachomatis* was prepared as previously described (Dumoux, 2012). HeLa cells were cultured on grids and 24 h after seeding infected with *C.trachomatis* LGV2 as previously described (Dumoux, 2012). 24 h post infection cells were plunge frozen using a Vitrobot (Thermo Fisher Scientific). Grids were pre-incubated on the grid with media containing 10% glycerol.

#### R. Rubrum

*Rhodospirillum rubrum* (gift by D. Canniffe from the University of Liverpool) was grown photosynthetically on modified *Rhodospirillaceae* (DSMZ 27) medium as previously described (García-Sánchez et al., 2018). Bacteria were exposed to 100 μmol photons/m^-2^/s^-1^ illumination at 30 °C in 150 ml flasks until an A680 of 1.6. 50 ml of culture was harvested by centrifugation at 3000 g for 30 min then resuspended with 15 ml of 20 mM Tris buffer at pH 7 and used for grid preparation.

8 μl of sample was applied to a plasma cleaned (Harrick plasma cleaner, medium setting, 40 s) Quantifoil Cu 300 R2/2 grid and plunge frozen in liquid ethane with a GP2 (Leica Microsystems, Wetzlar, Germany) blotted from behind, 6 s blotting time; 15°C and 75 % chamber temperature and humidity, respectively.

#### Brain tissue

68 day old Protamine-EGFP (PRM1-EGFP) mice (Haueter et al., 2010) (CD1; B6D2-Tg ^(Prm1-EGFP)^#Ltku/H) were euthanised in a schedule 1 procedure via intraperitoneal injection of sodium pentobarbital followed by decapitation following licensed procedures approved by the Mary Lyon Centre and the Home Office UK. Brains were dissected and cut into four equidistant lateral sections using a scalpel, with region one encompassing the olfactory bulb. Regions 2 and 3 were then further sectioned using a vibratome (VT1000S, Leica) set to produce 200 μm sections. Brain sections were kept at 4°C in Hank’s Balanced Salt Solution (HBSS) from death to high-pressure freezing. A 2 mm biopsy punch was used to excise regions of cortex from lateral slices.

Punches were then placed onto electron microscope grids during freezing. Brain punches on grids were frozen between 3 mm planchettes/carriers (Science Services, Munich, Germany) (assembled on mid-plates. Briefly, planchettes were coated with 1% soya-lecithin dissolved in chloroform and the solution allowed to evaporate. This process generates small microvesicles on the surface of the planchette. A flat-sided planchette was placed flat side upwards and a glow discharged (Glocube, Quorum, Lewes, UK) electron microscopy grids (UltraAuFoil, 200 Au mesh, 2/2 Au film (Quantifoil Micro Tools) placed on top; brain punches were then placed onto the grid and submerged in 20% bovine serum albumin in HBSS. 3 mm planchettes with a 0.1- or 0.2-mm recess, also coated with 1% soya lecithin, were then placed recess-side-down onto the assembled sandwich and high pressure frozen in a Leica HPM 100 (Leica Microsystems). The planchette grids were then disassembled under liquid nitrogen, clipped into Autogrids (Thermo Fisher Scientific) and stored at 80 K (liquid nitrogen) for later use.

#### Heart tissue

An 8-day old ‘wildtype’ C56BL/J mouse was euthanised in a schedule 1 procedure via intraperitoneal injection of sodium pentobarbital followed by decapitation following licensed procedures approved by the Mary Lyon Centre and the Home Office UK as described for the brain tissue. Heart was dissected and placed in ice cold Millonig’s buffer (Fisher Scientific, ThermoFisher Scientific) then fixed using ice cold 4% PFA in Millonig’s buffer (Fisher Scientific) then left to incubate overnight at 4°. Thin sections of 50μm were obtained using a vibratome and placed onto a glow discharged electron microscopy grid (UltrAuFoil, 200 Au mesh, 2/2 Au film (Quantifoil Micro Tools)) that had been pre-clipped into Autogrids (Thermo Fisher Scientific). The sample was assembled in the mid-plate between two 6mm planchettes incubated with hexadecane for 15 min. 20% w/v bovine serum albumin (BSA) (Sigma-Aldrich, St. Louis, Mo, USA) in PBS buffer was used as a cryoprotectant and filler and the assembly was high pressure frozen in a Leica HPM 100 (Leica Microsystems). The planchette-autogrid assembly was disassembled and stored under liquid nitrogen.

#### Yeast

*S. cerevisiae* were grown to an OD of 0.8 and plunge frozen as described (Khavnekar et al., 2022).

#### Beads

PEG 300 coated 1 μm polystyrene particles (Abvigen, Newark, NJ, USA) were diluted to 4 mg/ml and mixed 1:1 with 3x diluted 6 nm BSA-Gold nanoparticles (Aurion, Wageingen, Netherlands). 3 μl of the resulting mixture was spotted onto glow-discharged copper EM grids (Quantifoil 300 Au mesh, R2/2 carbon film (Quantifoil Micro Tools). The grids were then blotted for 2 or 4 seconds (blot force −10, humidity 70%) before plunge freezing by rapid immersion in liquid ethane using the Vitrobot (Thermo Fisher Scientific). Grids were subsequently clipped.

### FIB SEM

The serial plasma FIB/SEM imaging has been performed on a dual beam FIB/SEM “Helios G4 Hydra” with equipped with an Aquilos II type cryo-stage, a four source “Hydra” plasma ion column and an “Elstar” SEM column (Thermo Fisher Scientific). The detectors are an Everhart-Thornley Detector (ETD) and a through-the-lens detector (TLD) using a positive bias (70 V) suction tube. An immersion field is generated to improve detection and signal to noise. The samples were loaded into a custom purpose 25° pre-tilt holder. During the experiments, an instrument defect which required the development of a new ion gun design affected plasma ion beam capability, preventing the usage of argon at 30 kV. All experimental dataset using argon as source for milling were performed at 20 kV. We were able to produce a complete set of curtaining scores for all gas at 20 kV and 30 kV after the source had been successfully replaced.

#### Focused ion beam (FIB) milling

The accelerating voltage used for ion beam milling was 30kV, except for argon which was used at 20 kV or 30 kV.

To protect the leading edge while milling and to reduce surface topography, an organic platinum layer (trimethyl(methylcyclopentadienyl)platinum(IV)) of ~1-2 μm was deposited on the sample using a gas injection system (GIS). To reduce charging the sample was coated with two sputter coated layers (before and after the organic platinum GIS layer) of platinum ions using an in-built platinum mass as a sputter target; removal of platinum from the mass deposits platinum vapour onto the sample. The mass is placed at the end of a small rod that can be inserted into the path of the ion beam (16 kV and ~1 mA, with full view of the rod). Thickness of the sputtered layer depends on the exposure time of the target to the ion beam. We usually deposit each layer for 45 s which gives a layer that is tens of nanometres in thickness.

Once an area of interest has been identified, the sample is placed at eucentric height, and a first opening trench is milled at ~2 nA. This enables a check that the sample of interest is in the milled area and assess the sample quality. The surface is then polished before starting the milling and imaging sequence.

#### SEM

SEM image parameters are shown in the following table. Image acquisition took between 2 and 5 minutes. Parameters for the dataset are as below.

The electron dose has been calculated using this formula ED=(N/A) x t, where ED is the electron dose in e^-^/nm^2^, N is the number of electrons, A is the area and t the total time the sample has been exposed. To calculate N, we used N=I/e, where I is the current measured using a Faraday cup and e is the charge of an electron.

#### Serial FIB

The SerialFIB software (Klumpe et al., 2021) was run on the microscope via the AutoScript 4 (ThermoFisher Scientific) interface. In order to extend SerialFIB to plasma ion sources, the template pattern files that follow the Thermo Fisher standard were adjusted to a plasma FIB equivalent to allow reading and writing of patterns. Parameters that changed in those files between a system utilising gallium, i.e., an Aquilos 2 dual-beam FIB/SEM (ThermoFisher Scientific), to the system used in the current study utilising plasma-based milling were: total diameter, volume per dose, total beam area, sputter rates and depth per pass. One crucial parameter that had to be adjusted when switch ion sources was the milling pattern pitch, as the pitch in plasma-based ion sources needs to be significantly higher than when milling with gallium based liquid metal ion sources, likely due to the decrease in focussing capability of the plasma FIB. The pitch values were 9.5 nm for gallium and 116 nm for nitrogen and argon-based milling. Furthermore, the beam shift limits for the ion and electron column had to be adjusted to the limits on the Helios Hydra system.

The imaging script that allows for definition of a milling and another imaging position to allow for stage movements between the two steps in the volume imaging protocol was developed inside SerialFIB’s scripting interface. In brief, the function allowing for serial FIB – SEM volume imaging runs was modified to take one stage position for slicing and one stage position for imaging into account. While stage movements normally lead to adjustment to the SEM focus according to linked stage height, the focus was enforced to a user defined or via autofocussed determined value to remove the necessity of autofocussing after every stage movement. In addition, the horizontal field width was locked to avoid slight changes when readjusting focus after stage movements to avoid changes in the pixel size.

The script and all code for operating the microscope utilised here is available on the SerialFIB GitHub https://github.com/sklumpe/SerialFIB/.

#### AutoSlice and View (Thermofisher Scientific)

AutoSlice and View (ASV) version 4.2 was run on the Helios G4 Hydra. A fiducial was manually milled on the side of the target to assist the drift correction during milling. The fiducial should have a ‘Uniqueness score’ determined by the software or minimally 70%. No Y-correction was applied. On the preparation tab, only the ‘Green clean pattern’ was used to polish the surface (5-10 slices with an overlap of 5-10). Regarding the imaging parameters, no alignment, Y-shift correction, auto-focus, auto source tilt, auto contrast and brightness was used, and we did not use tiling.

### Image metrics

#### Curtaining propensity

Plunged-frozen *C. trachomatis* infected HeLa cells were coated with a protective organoplatinum layer and platinum sputter coated as described. Each plasma ion beam was aligned immediately prior to milling. To enable effective curtaining rate determination with nitrogen and oxygen, double spot compensation (Figure 1 - supplementary Figure 4) was performed for each current.

As the practical usable current is determined by the apertures within the ion column, the current used for each gas for study is based on the use of the same aperture; for example, at the lowest aperture size used the currents for nitrogen, oxygen, xenon and argon are 0.27 nA, 0.23 nA, and 0.1 and 0.2 nA, respectively. A list of currents for a given aperture is shown in Table 2. However, these values are the ones displayed and will change over time with the beam milling its own apertures.

**Table 1:** serial pFIB/SEM parameters used for the different biological samples.

| sample | milling gas | acceleration voltage (kV) | target current (nA) | software | SEM imaging angle (deg) | pixel size (nm) | voltage SEM (kV) | current SEM (pA) | Dwell time (ns) | Line integration | gain | voltage offset (V) | electron dose (e/A2) * |
| --- | --- | --- | --- | --- | --- | --- | --- | --- | --- | --- | --- | --- | --- |
| HeLa | argon | 20 | 0.2 | TFS ASV | 52 | 6.745 | 1.25 | 6.25 | 100 | 100 | 62.9 | -12 | 8.24E-04 |
| Vero | nitrogen | 30 | 0.27 | Serial FIB | 90 | 1.927 | 1.2 | 6.25 | 100 | 100 | 56.45 | -8.4 | 9.84E-03 |
| *R. Rubrum* | argon | 20 | 0.2 | Serial FIB | 90 | 1.942 | 1.1 | 6.25 | 100 | 100 | 36 | -3.37 | 9.94E-03 |
| brain | argon | 20 | 0.2 | TFS ASV | 52 | 6.745 | 1.2 | 6.25 | 500 | 16 | 40.25 | -4.37 | 8.24E-04 |
| heart | argon | 20 | 0.2 | TFS ASV | 90 | 5.62 | 1.1 | 6.25 | 100 | 100 | 45.42 | -4.41 | 1.19E-04 |
| * formula from ... current has been measured using a faraday cup |  |  |  |  |  |  |  |  |  |  |  |  |  |

**Table 2:** FIB current (targeted value) in nA.

|  | Argon | Xenon | Nitrogen | Oxygen |
| --- | --- | --- | --- | --- |
| 20kV | 6.2 | 3.3 | 14 | 4.2 |
|  | 2.4 | 1 | 1.8 | 1.4 |
|  | 0.2 | 0.1 | 0.32 | 0.29 |
| 30kV | 7.6 | 4 | 24 | 5.6 |
|  | 2 | 1 | 1.7 | 1.7 |
|  | 0.2 | 0.1 | 0.27 | 0.23 |

A series of 5 windows of 2 x 2.5 x 2 μm were milled until the average grey level intensity for the window reached 10% of the initial signal. The acceleration voltage for the I-beam was kept at 30kV or 20kV. To accurately reflect the ion beam current applied on the sample, the current presented for this experiment is the measured current. The milled surface is then imaged 90° to the SEM without an electrostatic lens to avoid interaction of the magnetic field with the ion beam. This is crucial for imaging with oxygen and nitrogen, where the use of the pre-field immersion lens led to blurring of the ion beam images. The pixel size, voltage, current, electric gain, and voltage offset were kept identical. This was repeated three times per current (total of 15 windows per current).

As curtaining can vary significantly across a milled surface, an automated, quantitative assessment of curtaining is necessary for un-biased comparison of ion sources and milling settings. We use a method based on the work from Hovden’s group (Schwartz et al., 2019). Using Fiji (ref) the initial image is subject to a fast Fourrier transform (FFT). A horizontal 5° mask is applied and a reverse FFT performed. In this new image the vertical lines have been removed. By subtracting the initial and inverted masked FFT images, we can produce a mask to isolate those lines using a default threshold based on Minimum Cross Entropy (Li & Tam, 1998). A merged image where this mask is superimposed to the initial image allows precise identification of the location where the milling was performed. We can then produce an histogram where the white pixels are part of the curtain. The percentage of this white pixels is calculated and forms the curtaining score (Figure 1 - supplementary figure 1).

#### FRC calculation

The FRC calculation for measuring local resolution was derived from the method of Koho *et al* (Koho et al., 2019), which was originally developed for optical microscopy. Their one-image FRC calculation has a calibration factor applied, which was derived from optical microscopy images. This calibration aims to match the one-image FRC resolution value to the gold-standard two-image FRC value. To recalibrate this measure to SEM images, their calibration procedure was repeated (Figure 4 - supplementary figure 2).

Here, the one-image FRC was used to measure local resolution as a quantitative comparison between images at different acquisition settings and positions within the stack. The images to be studied here were split into patches of 256 x 256 pixels, and the image resolution for each patch was calculated with the newly calibrated one-image FRC measure.

To calibrate the one-image FRC for EM images, a calibration dataset was obtained which consisted of pairs of images taken of the same FOV at different pixel sizes (1.12, 2.25, and 4.5 nm/pixel). One set of images was taken at 52° SEM angle, and the other at 90°. Both images in each pair were registered to each other with SIFT landmark placement using the linear stack alignment with SIFT plugin in Fiji 2.3.0/1.53q (Schindelin et al., 2012). The images were cropped to one FOV for all pixel sizes for each SEM angle, which covered 2000 x 2000 nm and 1600 x 1600 nm for the 52° and 90° images respectively, note that the number of pixels in each pair of images was different due to the varying pixel size.

The two-image FRC curves were calculated from each pair of images at each pixel size and SEM angle. Briefly, the cross-correlation between both images in the Fourier domain within frequency bands was determined, and the cross-correlation values were plotted against the frequency. The frequency was normalised against the maximum frequency values for each image to enable direct comparison between images. The normalised frequency at which the cross-correlation falls below 0.143 was determined, which is represented as *r_ref_* for the two-image FRC, and *r_co1_* for the uncalibrated one-image FRC. The one-image FRC curves were obtained for each of the images in the pair using the checkerboard sampling approach described in the work of Koho *et al*.

A calibration curve was fitted to the plot of 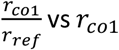 vs *r_co1_* (Figure 4 - supplementary figure 2). Application of this calibration factor to the one-image FRC curve shifted the curve to match the gold-standard two-image FRC measure, so the resolution value from the one-image FRC was comparable to the reference measurement (Figure 4 - supplementary figure 2). The original implementation of Koho *et al*. was modified to include this calibration so that it would be applicable to SEM images. This modified one-image FRC code is available as part of Quoll, an open-source Python package (Ho, 2022).

#### Shift measurement

1 μm PEG beads samples were used (see previous section). Argon 30 kV 200 pA was used. SEM images were acquired using 1.2 kV 6.25 pA exposing for 100 ms. In order to reduce the charging artefact, we use line integration of 25.

To adequately calculate the extent to which the samples shift during tilting, SIFT landmark placement in Fiji/ImageJ 2.3.0/1.53q was used to automatically place matching landmarks between pairs of adjacent Z slices. The Euclidean distance was calculated between each pair of landmarks with the formula:

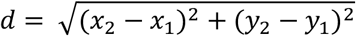

The Euclidean distance between landmarks was divided by the diagonal length of the FOV of the image, to ensure a fair comparison between images of different sizes.

#### Depth of field

A tin ball sample was used and, with the focus fixed, the stage height was adjusted in ~5μm increments to obtain a through-focus series of images. The FRC method was applied to each of these to estimate the average resolution, giving a clear minimum (Figure 4 - supplementary figure 5). The depth of field was then taken as the distance through which no significant change in the image resolution was measured. This was carried out by a two-tailed, two sample equal variance t-test was carried out with a confidence interval of 95% between the measured resolutions at each stage position with the stage position with the lowest measured resolutions. For both 1 kV and 2 kV, four stage positions were found to be statistically similar (marked with asterisks in Figure 4 - supplementary figure 5), so the depth of field was therefore estimated to be ~20μm.

#### Sphericity (3D)

From the images obtained, an average size of the beads can be extracted and compared with manufacturer values. Assuming the beads are perfectly spherical, the images are expected to show a cross-section of the bead as being circular. Mathematically, the area covered by the bead is expected to be consistent with the area of a circle with a varying radius along the perpendicular slice direction. This can be derived from the sphere equation, and the area is equal to 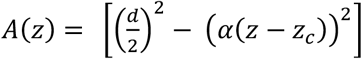, with *d* being the diameter of the bead, *z* the slice coordinate, and *z_c_* the height centre position of the bead. The parameter *α* was added here to consider potential coefficient error in the milling thickness, which in a perfectly calibrated milling-capable microscope would be exactly 1. The experimental slice-thickness can be estimated by multiplying *α* with the target milling thickness. In each image slice where beads were annotated using plugin for Napari pyclesperanto

(Sofroniew et al., 2022) (https://github.com/clEsperanto/pyclesperanto_prototype), the two-dimensional area of each bead was collected from the number of pixels, and plotted as a function of slice as dots, for each of the methods used. The respective lines represent optimised *A*(*z*) [https://docs.scipy.org/doc/scipy/reference/generated/scipy.optimize.curve_fit.html] curves, with parameters *d, z_c_* and *α* being allowed to vary for each of the beads independently

#### Circularity (2D) (Chlamydia infected cells)

Bacteria were manually segmented, and the outline analysed using the ‘Analyze Particles’ tools from FIJI (Schindelin et al., 2012). The average and standard deviation were then calculated and shown in Figure 4 - supplementary figure 1.

#### Charging artefact mitigation

Segmentation of charging centres was completed using a FastAI version 1.06 (Howard & Gugger, 2020) implementation of a 2D U-Net (Ronneberger et al., 2015). The U-net was initially set up with an encoder based in resnet34 and training was done on 256px x 256px patches of 1024 test images and respective manually segmented labels, with and without artefacts at a ratio of 1:1. Then this data was split into training (80%) and validation (20%) data. The preprocessing transforms used were FastAI’s package defaults and with imagenet normalisation. The loss function used was binary cross entropy (bce) applied to the logits, and weight decay was set initially to 0.01. A total of 10 epochs were run resulting in IoU score in the validation data of 0.90. This model was then used to automatically segment the charging artefacts of the whole data volume. Manual correction of this segmentation was undertaken and then compared using Dice coefficient (Dice, 1945) to the automated segmentation. The segmented and manually adjusted dataset was used for subsequent work. After segmentation, local areas near charging centres were assessed on a row-by-row basis, both from the top and bottom of the image. Areas with charging artefacts were then filtered using the average of the previous 20 rows. The following smoothly varying continuous functions were then fitted and subtracted from the filtered data line: (https://github.com/rosalindfranklininstitute/chafer).

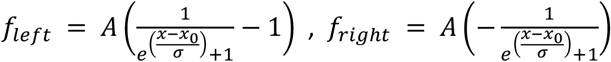

with *A, x_0_* and *σ* being parameters to be fitted, and functions *f*_left_ fitted on the left of the labelled artefact, and *f_right_* fitted on the right side. Other smoothing functions (gauss tail, exponential, 1/x) were tested but these gave far better overall results both in fitting to the shape of the horizontal tails and algorithmically successful in finding optimal solutions.

In total, 117 charge centres were identified during segmentation of the *S. cerevisiae* dataset. At each of these charging centres, an average line profile of the grey values in both the X and Y directions was taken. For the horizontal line profiles, the box over which the line profiles were averaged was 1x the charging centre diameter in height and 9x in width, while for the vertical line profiles, it was 1x in width and 9x in height. The box size was chosen empirically to capture the full extent of the charging artefact. Due to the presence of other charge centres nearby or to the edges of the image, 20 charging centres were assessed fully.

#### Movies

Raw data have been deposited in the EMPIAR data repository (XXXX). All movies have been aligned using SIFT (Lowe, 2004) and filtered using a mean (2 pixels) filter using either Fiji/ImageJ 2.3.0/1.53q or Amira 2020.2. The data acquisition parameters of the SEM are described in the FIB/SEM section.

#### Segmentation and quantification

Amira Version 2020.2 (ThermoFisher Scientific) was used to align and segment the data set. The alignment was performed using the DualBeamWizard and segmentation was carried out manually. For volume/surface rendering, the object corresponding to each segmentation was exported as tiff stacks and visualised in 3D viewer in fiji (Schindelin et al., 2012). For the quantification, the object containing the manually segmented boundary was exported from Amira as tiff stacks. These stacks were converted to binary 3D masks and ImageJ 3D object counter was run on the region of interest. Results exported as a .csv document for analysis in Excel.

## Acknowledgements

The authors would like to thank the experimental coordinators at Diamond Light Source for their diligence in supporting our instruments, including during the overnight runs. We would like to thank Chelsea Norman for her support in sample preparation, and Chloe Cheng for helpful discussion and feedback on the image processing workflows. We would like to thank Dr Silvia da Graça Ramos and Despoina Eugenia Kiousi for their help with segmentation. We would like to thank D. Canniffe from the University of Liverpool for providing *R. rubrum* samples. We would like to thank Marianne Yon, Alexandra Rodrigues and Sara Wells of the Mary Lyon Centre at MRC Harwell for their support with animal work. The Rosalind Franklin Institute is funded by UK Research and Innovation through the Engineering and Physical Sciences Research Council. Funding was also provided by the Wellcome Trust through the Electrifying Life Science grant (220526/Z/20/Z to JHN).

## Authors contribution

Sample preparation: MD, PYAL, WB, JLRS,DF, LW

Imaging MD, TG, MG, JLRS

Data processing MD, DF, JLRS, MG, EMLH, NBY, LMAP, MB, MCD

Designing experimental procedure MD, TG, MCD, JHN, MG, RK

Designing software MCD, MB, SV, SK, JMP

Software implementation EMLH, NBY, SK, LMAP, MB

Writing paper MD, EMLH, NBY, SK, LMAP, DF, DKC, CAS, MCD, JHN, MG

Reviewing paper MD, TG, EMLH, NBY, SK, LMAP, RK, DF, PYAL, WB, JLRS, JP, LW, MB, DKC, CAS, MCD, JHN, MG

## Data and code availability

Raw data are deposited on the EMPIAR data repository (XXXX). Code could be found on the Rosalind Franklin Institute GitHub (https://github.com/rosalindfranklininstitute/) and the serialFIB GitHub (https://github.com/sklumpe/SerialFIB/)

## Supplementary material

### Supplementary movies

Figure 2 - supplementary movie 1: volume from serial pFIB/SEM of HeLa cells after alignment, mean filter and cropping to the region of interest.

Figure 3 - supplementary movie 1: volume from serial pFIB/SEM of live slice of mouse brain after alignment, mean filter and cropping to the region of interest.

Figure 5 - supplementary movie 1: volume from serial pFIB/SEM of *R. rubrum* after alignment, mean filter and cropping to the region of interest.

Figure 6 - supplementary movie 1: volume from serial pFIB/SEM of Vero cells after alignment, mean filter and cropping to the region of interest.

Figure 6 - supplementary movie 2: segmentation (Amira) from the volume from serial pFIB/SEM of Vero cells after alignment, mean filter and cropping to the region of interest. Yellow: endoplasmic reticulum, red: lipid droplets, green: mitochondria, cyan: lipid droplets to ER membrane contact sites, purple: mitochondria to endoplasmic reticulum membrane contact sites.

Figure 7 - supplementary movie 1: volume from serial pFIB/SEM of fixed slice of mouse heart after alignment, mean filter and cropping to the region of interest.

### Supplementary Figures

**Figure 1 - supplementary figure 1:**
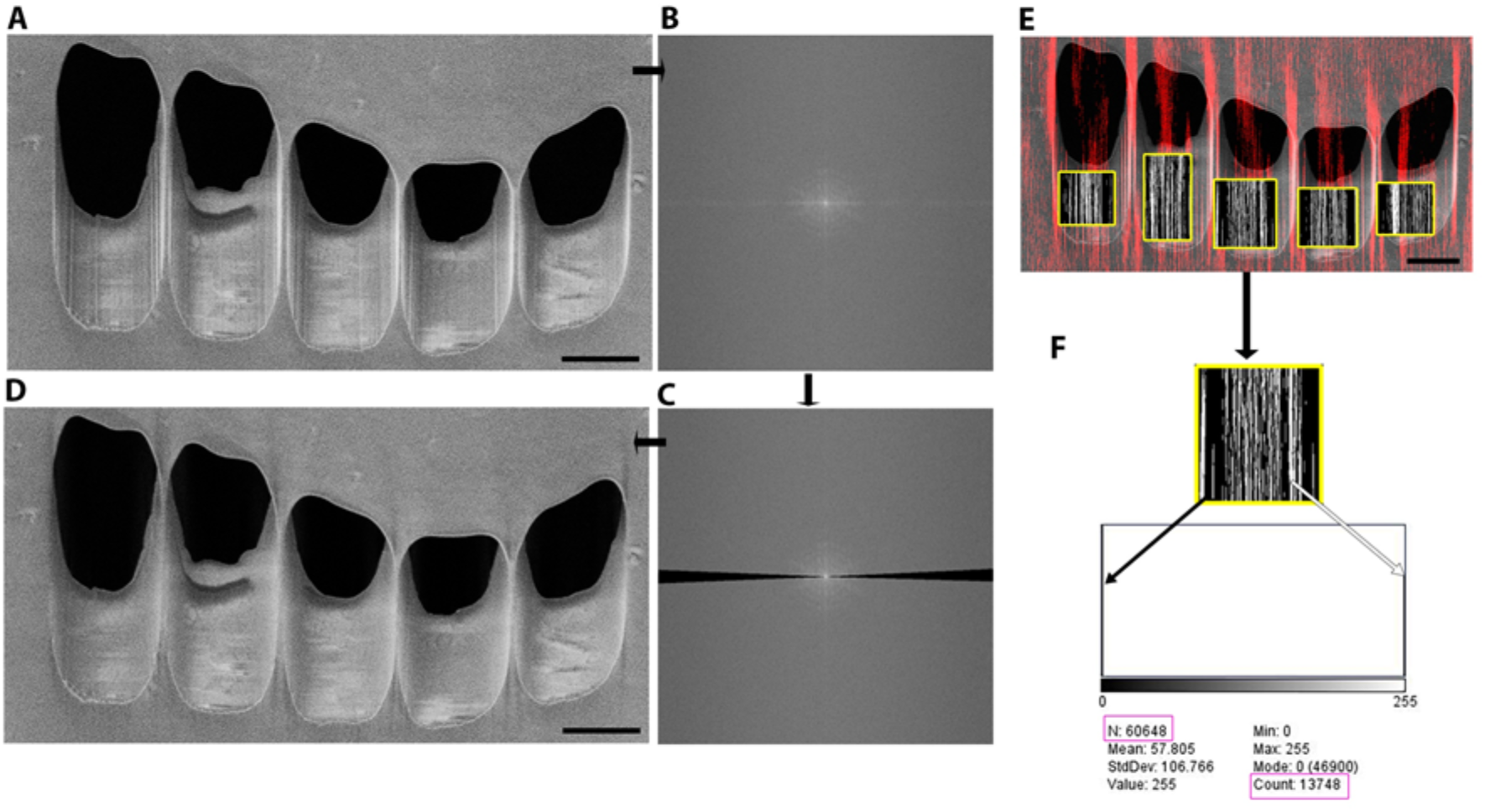
Methods to determine the curtaining score. (Material and Methods). Representative image analysis from HeLa cells were milled and imaged using the SEM at 90° incidence. In this example milling has been done using argon (30 kV,with measured current of 6.1nA). Using Fiji (Schindelin et al., 2012) initial image (A) with its corresponding FFT (B) has been filtered (C) producing a reverse FFT (D) where the vertical lines have been removed. Subtracting (D) from (A) we can produce a mask to isolate those lines and form a merged image where this mask (red/white) is superimposed to the initial image (E). From this we can precisely identify the location of the window which has been milled. This will allow to isolate the area within the mask and determine the number of pixels which are part of a curtain using the histogram (F) where white pixels are part of the curtain and their number appears in the count. The black pixels are not part of a curtain and the total number of pixels appears as N. Scale bar: Scale bar: 2μm.

**Figure 1 -supplementary figure 2:**
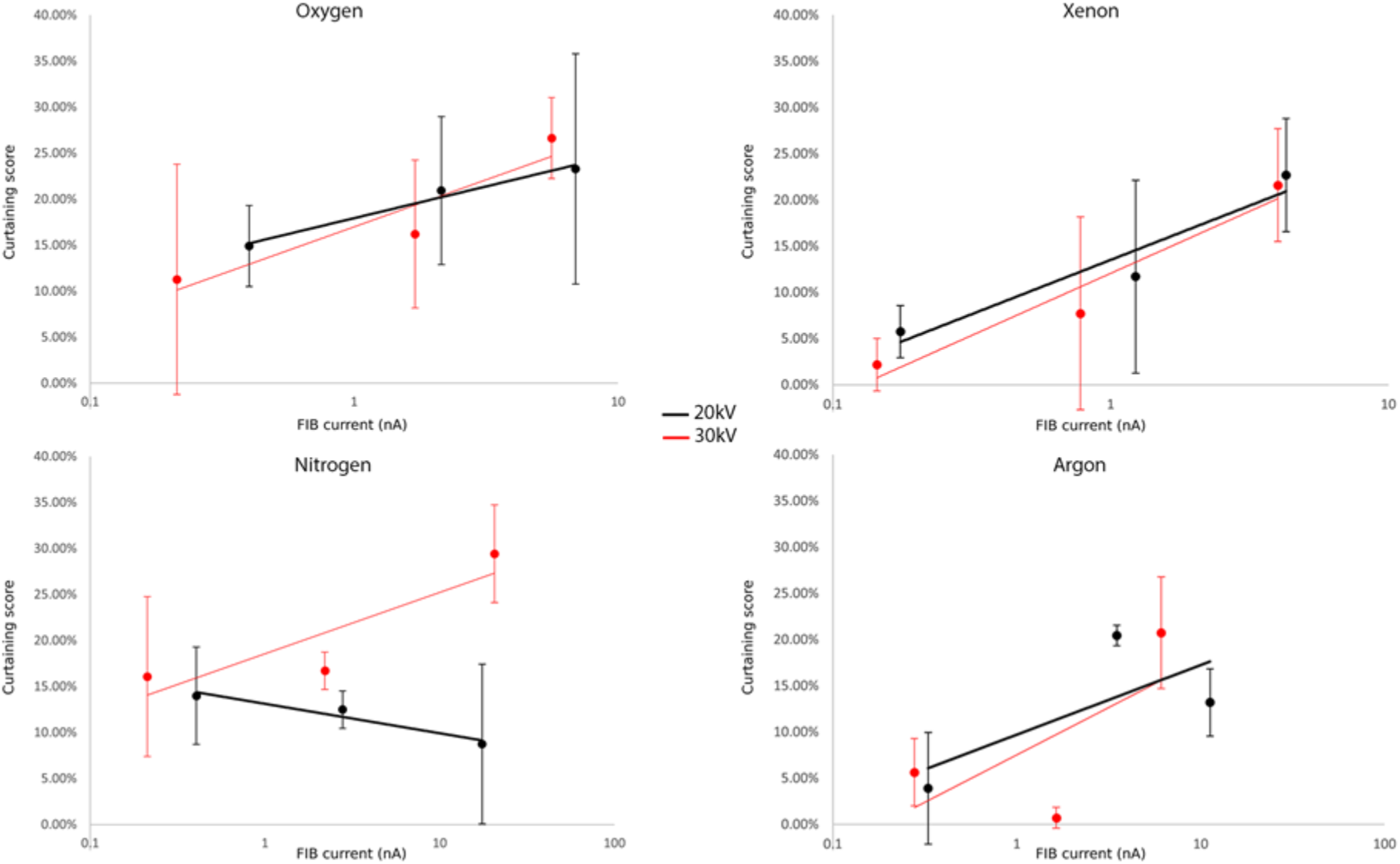
curtaining score for the different gases at 20 kV or 30 kV. Plunge frozen *C.trachomatis*-infected HeLa cells were milled for each current and each gas. 15 windows of 2×2.5×2μm were milled at different measured currents at the different acceleration voltage 30 kV (red) or 20 kV (black). The incidence angle was set to 18°. SEM images were acquired 90° to the FIB. Plot of curtaining score as a function of current (see materials and methods). Points represent the mean value associated with the standard error.

**Figure 1 - Supplementary Figure 3.**
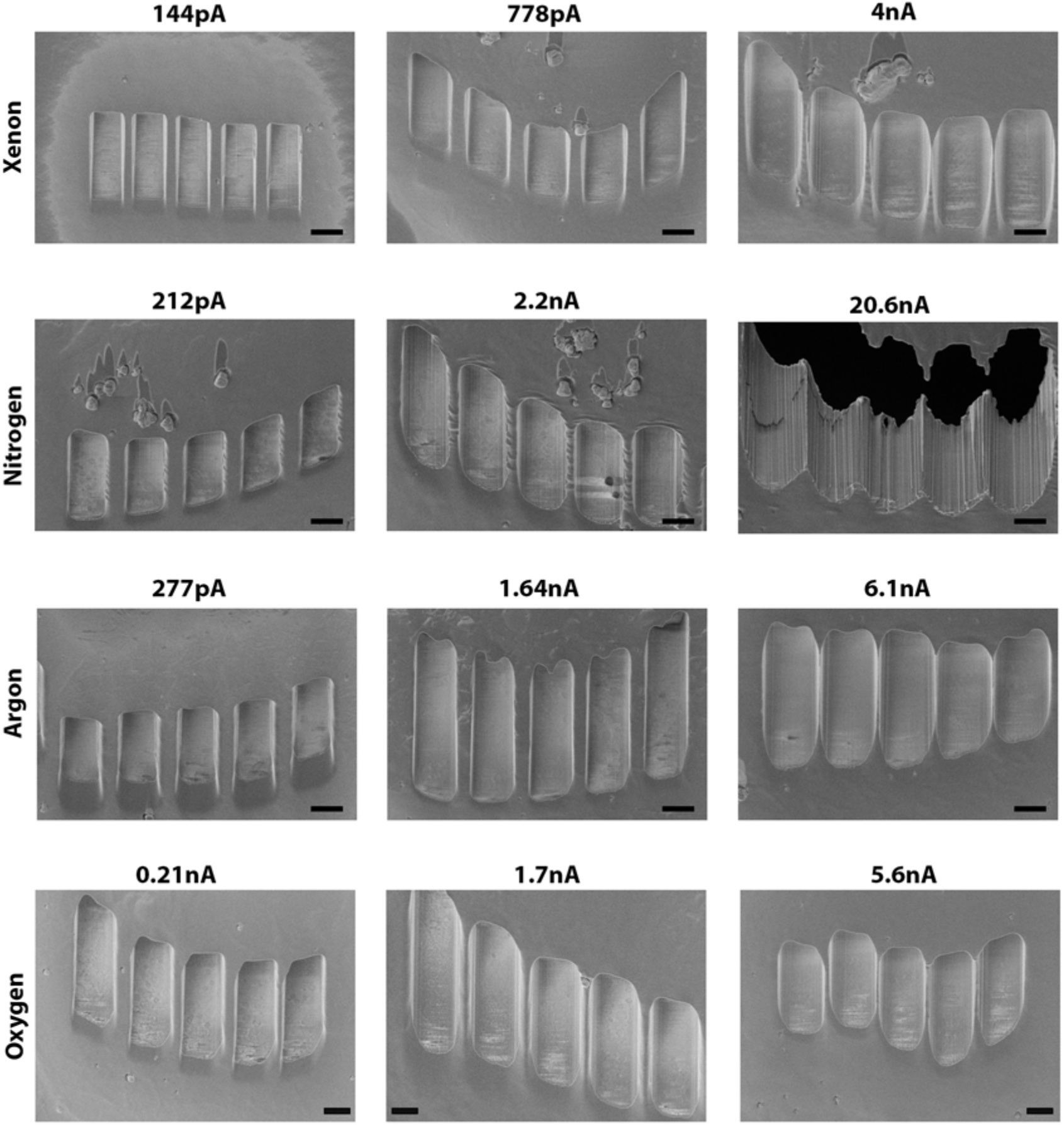
Example of curtaining effects obtained at different current for (top to bottom) xenon, nitrogen, argon, and oxygen, respectively. SEM images have been acquired at 90° incidence. Currents shown are the currents measured on the microscope during acquisition. Scale bar: 2μm.

**Figure 1 - Supplementary Figure 4.**
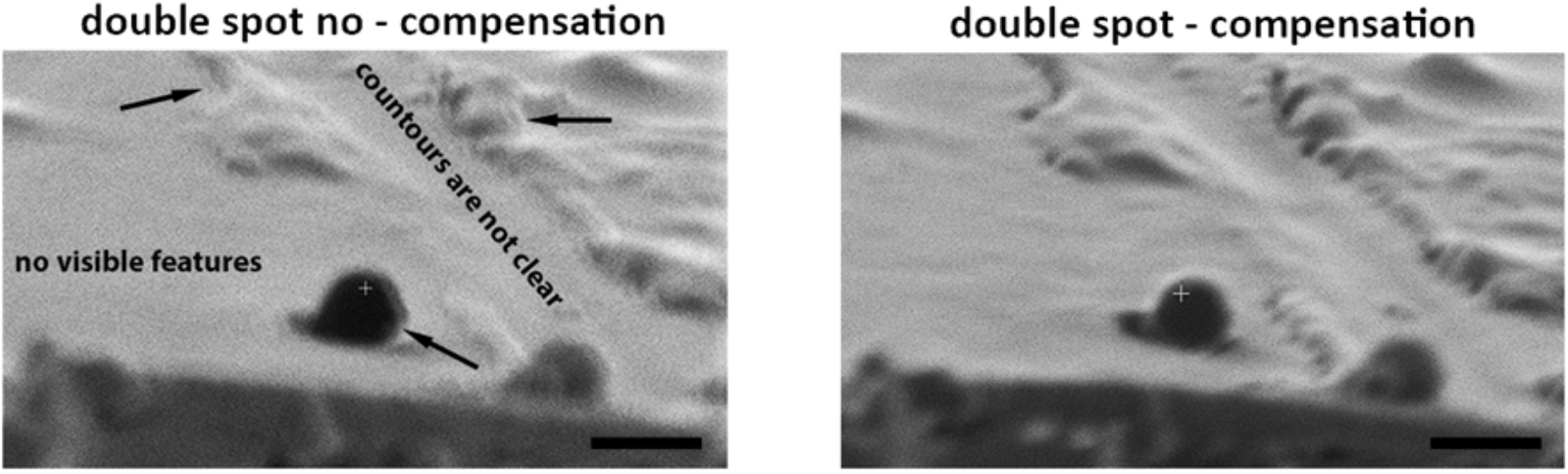
Example of FIB images acquired using Nitrogen as ion source with (left) or without (right) double image compensation. Arrows show where the lines are doubled. Overall contours are unclear and small features are not visible. Scale bars: 1μm.

**Figure 4 Supplementary Figure 1.**
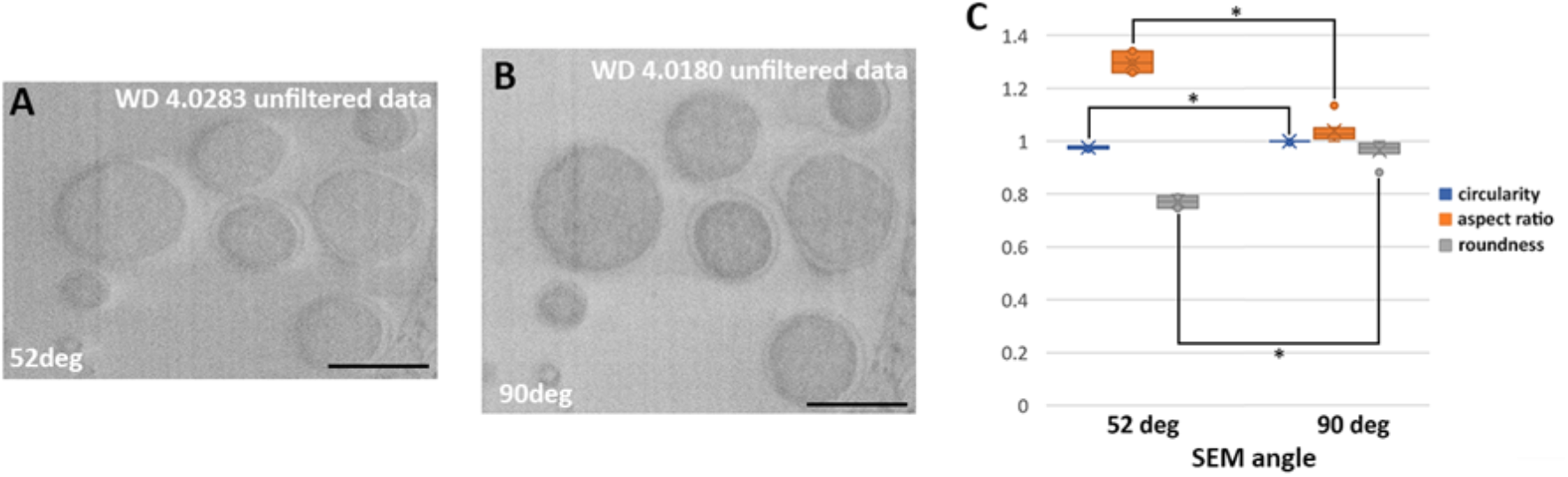
Effect of imaging angle on field of view. *C.trachomatis* infected HeLa cells were imaged using serial pFIB/SEM using argon to mill. (A) and (B) show images acquired at 52° and 90°, respectively. The same field of view of cells where bacteria can be identified was acquired. The angle is the only variable applied, with the working distance (WD) shown and all other imaging parameters kept identical. (C) Bacteria have been segmented and circularity, aspect ratio and roundness was calculated (Rueden et al., 2017) and presented in the graph in function of the imaging angle with the standard deviation. For reference, a sphere has a value of 1 for all three parameters. * indicates: significantly different (0.1%).

**Figure 4 – supplementary figure 2:**
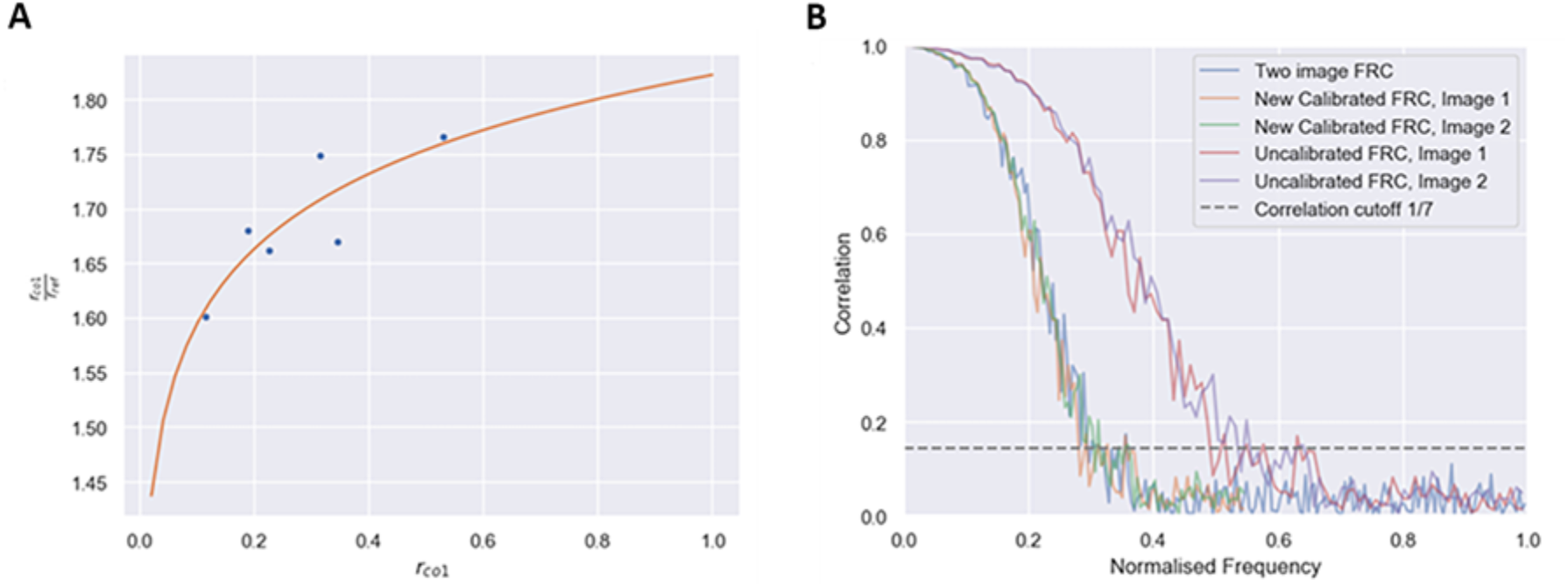
FRC evaluation method. (Material and Methods). (A) Calibration curve obtained from the EM calibration dataset. Each point represents a pair of images taken at each pixel size and each SEM angle. (B) Application of a calibration factor to the one-image FRC matches the FRC curve to the gold-standard two-image FRC, as shown by the orange and green lines (calibrated one-image FRC) closely following the blue line (gold-standard two-image FRC). The red and purple lines (uncalibrated FRC) do not match the gold standard FRC curve, so the resolution value taken from these curves are incorrect.

**Figure 4 Supplementary Figure 3.**
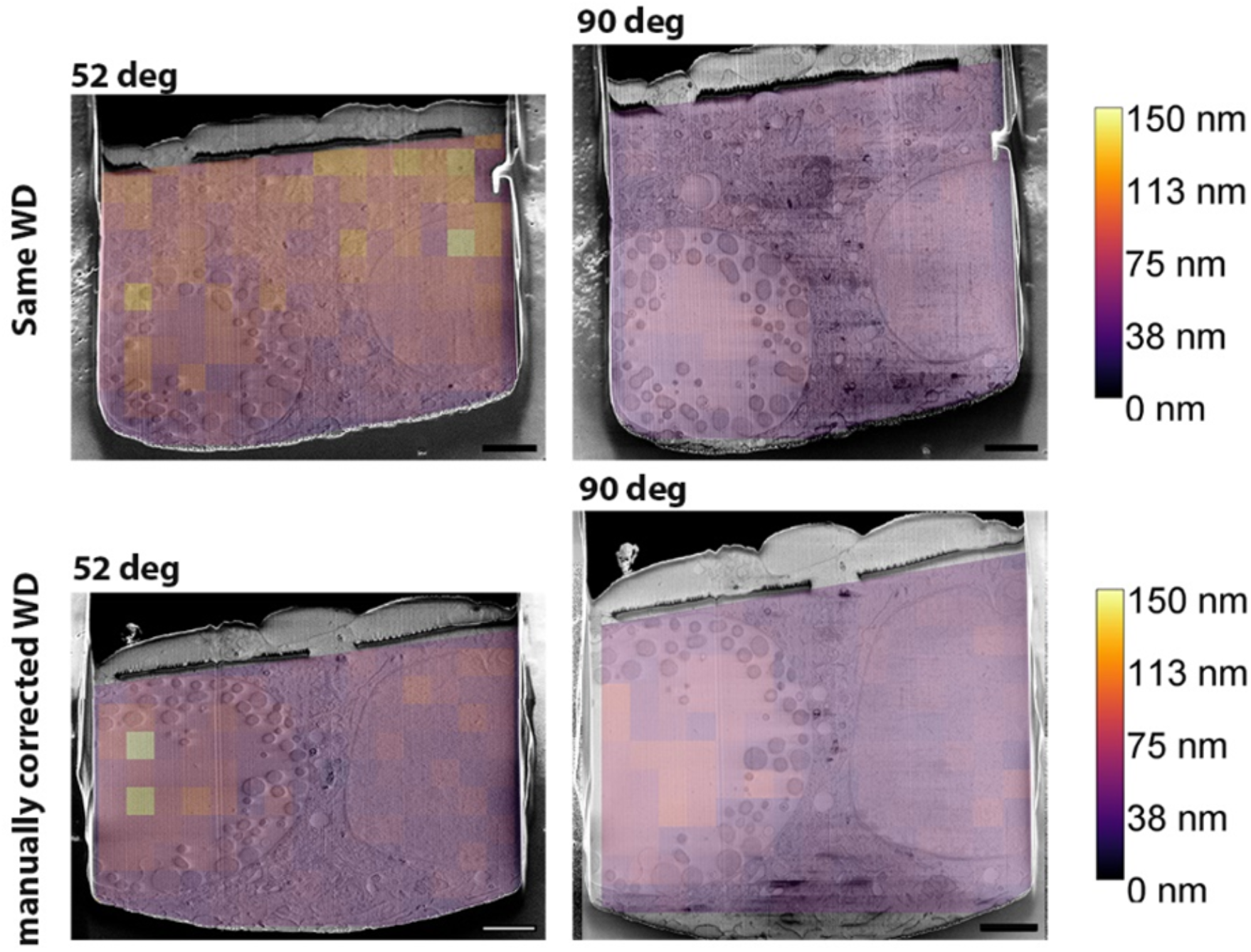
Relationship between resolution and working distance. SEM images taken at different imaging angles (52° or 90 deg to the SEM column) are overlaid with the results of the measured Fourier ring correlation resolution (heat map). Results are shown for instances where the working distance (WD) either remains the same (top) or not (bottom), while all the other parameters remain identical. Scale bar: 2μm.

**Figure 4 Supplementary Figure 4.**
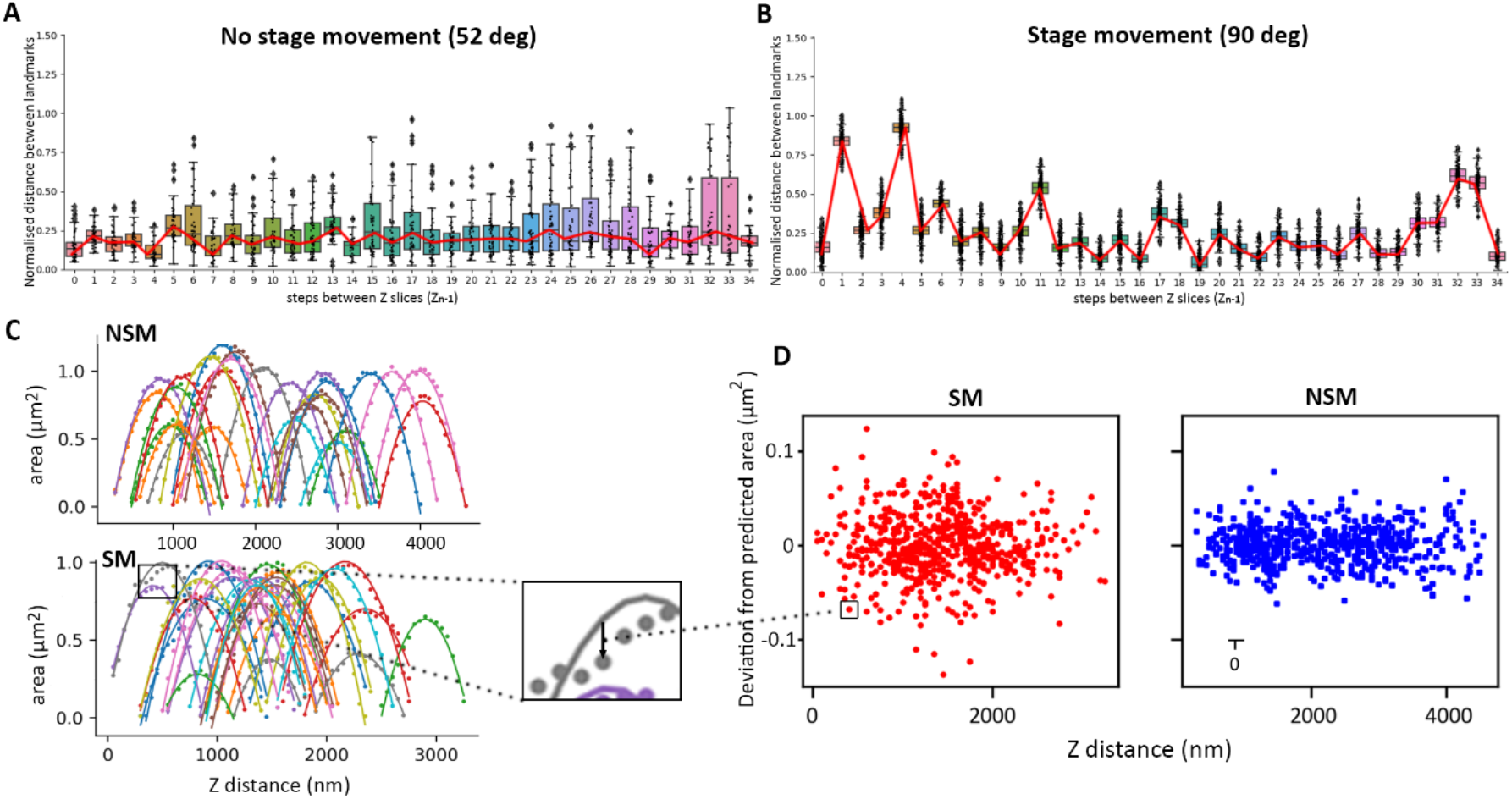
Measurement of XY (A and B) and Z drift (C and D) during data acquisition with no stage movement (NSM) imaged 52° to the SEM or with stage movement (SM), imaged 90° to the SEM. Serial pFIB/SEM stacks of one micron PEG beads were acquired with a 50 nm milling step using argon. (A and B) represent the normalised distance between bead landmarks (distance between landmarks/diagonal of the field of view) as a function of the step between slices (Z_n-1_). Red lines highlight the variation of the average in the position of the landmark (points). (C) Individual bead crosssection area as a function of Z position (Z distance in nm) which is the Z slice number multiplied by the target milling thickness (in this case 50nm). Points display the measured area from the dataset, and the solid line is the optimised curve for a perfect spherical bead. The deviation between prediction and measurement (example in the zoom box highlighted by an arrow) has been plotted (D) (deviation as a function of Z distance).

**Figure 4 Supplementary Figure 5.**
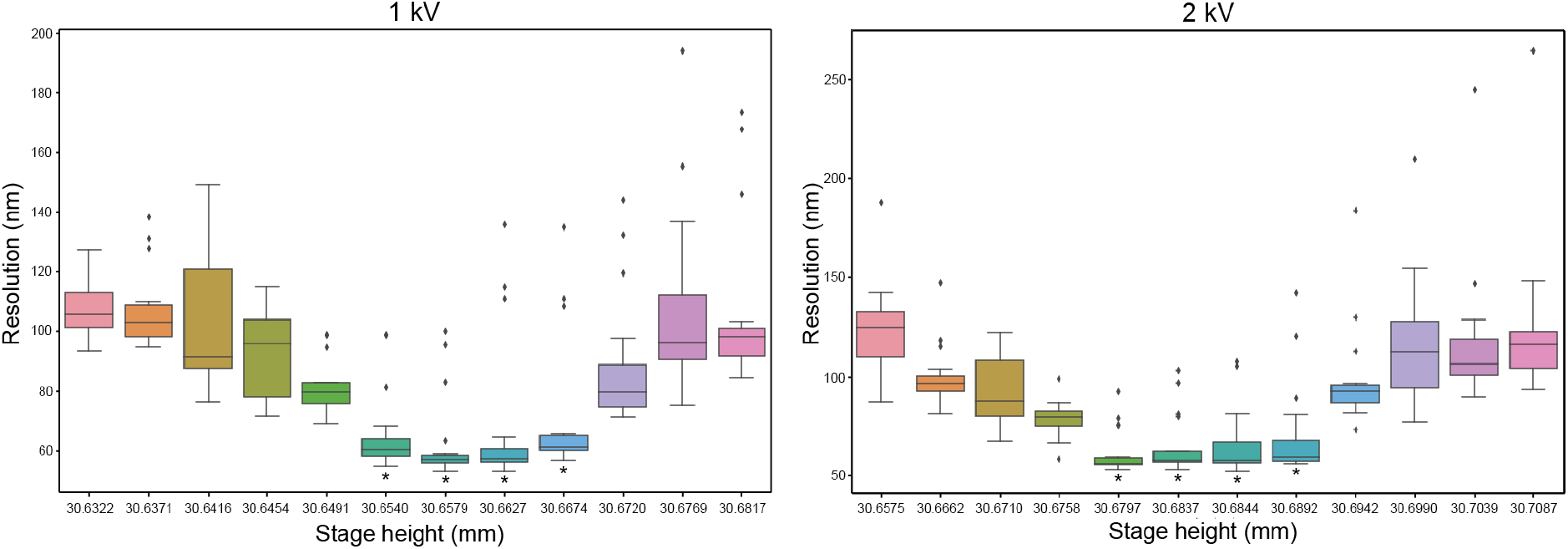
Measurement of depth of field. A tin ball sample was imaged at different stage position (Δ5μm) normal to the SEM using the immersion mode at 1 or 2 kV 12.5 pA. The FRC was then calculated and compared stage position by stage position to estimate if it was equal with 5% error (test). By taking the depth of field and measuring where the FRC is constant (T-test 95% confidence marked by an asterix), we can estimate the depth of field to be ~20 μm for imaging both at 1kV and 2kV.

**Figure 8 - Supplementary figure 1.**
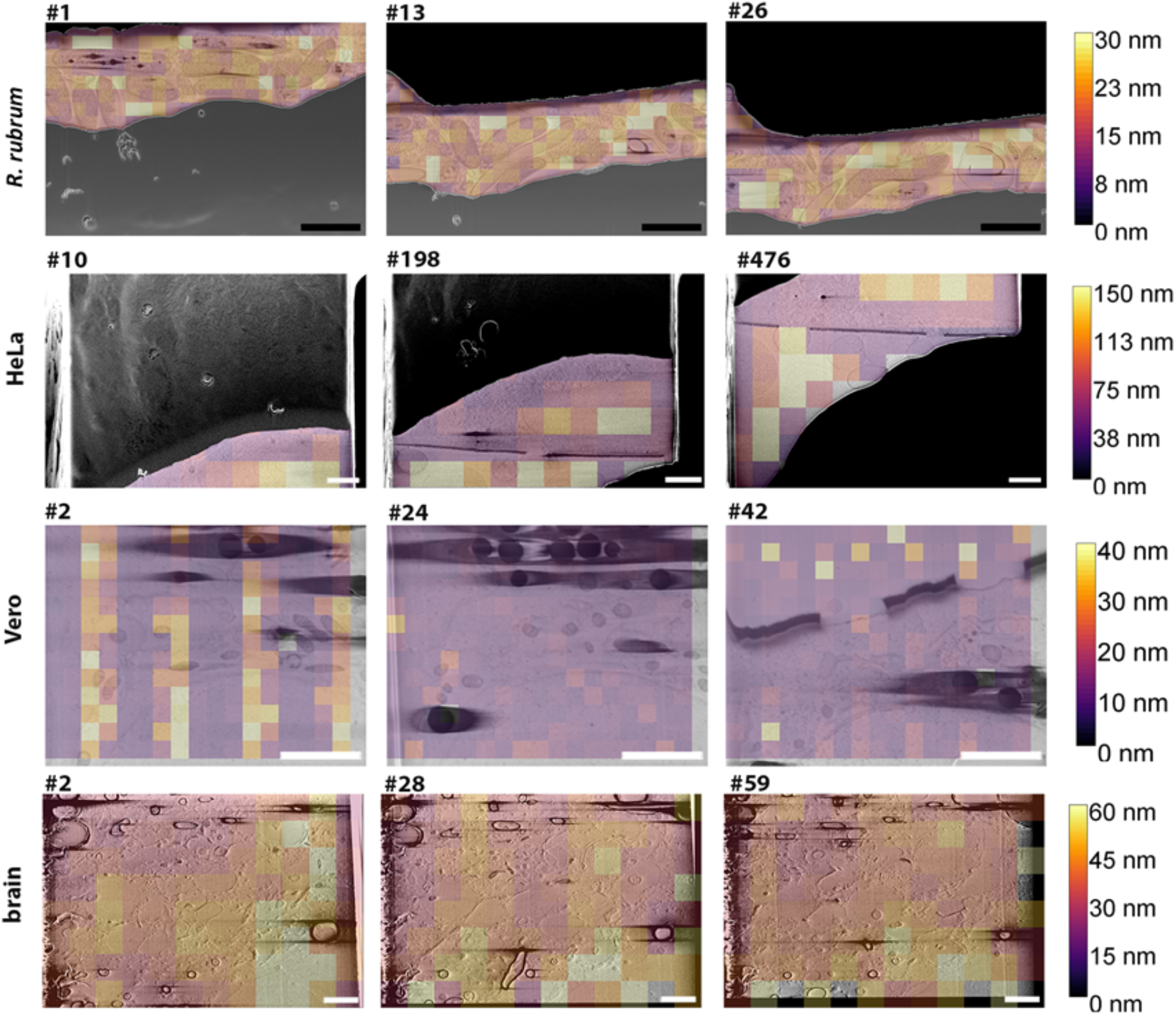
Analysis of the resolution from different biological data acquired. Slices from serial pFIB/SEM volumes taken at positions toward the beginning of the acquisition (left), at a point in the middle (middle) and towards the end (right). The slice number is shown above the respective images. The colour scheme relates to the locally measured resolution across the field of views using the Fourier ring correlation approach, are presented as overlays of the sample and the generated heat map (colour code on the right). Scale bars: 2μm

## References

Bäuerlein FJB, Baumeister W. (2021). Towards Visual Proteomics at High Resolution. Journal of Molecular Biology 433:167187. doi:10.1016/j.jmb.2021.167187

Begay RL, Graw SL, Sinagra G, Asimaki A, Rowland TJ, Slavov DB, Gowan K, Jones KL, Brun F, Merlo M, Miani D, Sweet M, Devaraj K, Wartchow EP, Gigli M, Puggia I, Salcedo EE, Garrity DM, Ambardekar A v, Buttrick P, Reece TB, Bristow MR, Saffitz JE, Mestroni L, Taylor MRG. (2018). Filamin C Truncation Mutations Are Associated With Arrhythmogenic Dilated Cardiomyopathy and Changes in the Cell–Cell Adhesion Structures. JACC: Clinical Electrophysiology 4:504–514. doi:10.1016/j.jacep.2017.12.003

Berger C, Dumoux M, Glen T, Yee NB y, Mitchels JM, Patáková Z, Naismith JH, Grange M. (2022). Plasma FIB milling for the determination of structures in situ. bioRxiv 2022.08.01.502333. doi:10.1101/2022.08.01.502333

Burnett, T. L., Kelley, R., Winiarski, B., Contreras, L., Daly, M., Gholinia, A., Burke, M. G., & Withers, P. J. (2016). Large volume serial section tomography by Xe Plasma FIB dual beam microscopy. Ultramicroscopy, 161, 119–129. https://doi.org/10.1016/j.ultramic.2015.11.001

Dice, L. R. (1945). Measures of the Amount of Ecologic Association Between Species. Ecology, 26(3), 297–302. https://doi.org/10.2307/1932409

DeRosier DJ. (2021). Where in the cell is my protein? Quarterly Reviews of Biophysics 54:1–32. doi:10.1017/S003358352100007X

Eder K, Bhatia V, Qu J, Leer B van, Dutka M, Cairney JM. (2021). A multi-ion plasma FIB study: Determining ion implantation depths of Xe, N, O and Ar in tungsten via atom probe tomography. Ultramicroscopy 228:113334. doi:10.1016/j.ultramic.2021.113334

Else, P. L., & Hulbert, A. J. (1985). An allometric comparison of the mitochondria of mammalian and reptilian tissues: The implications for the evolution of endothermy. Journal of Comparative Physiology B, 156(1), 3–11. doi.org/10.1007/BF00692920

Giacomello M, Pellegrini L. (2016). The coming of age of the mitochondria–ER contact: a matter of thickness. Cell Death & Differentiation 23:1417–1427. doi:10.1038/cdd.2016.52

Gorelick, S., Korneev, D., Handley, A., Gervinskas, Ge., Oorschot, V., Kaluza, O, Law, R. HP., Bryan, M. O., Pocock, R., Whisstock, J. C., & de Marco, A. (2018). Oxygen plasma focused ion beam scanning electron microscopy for biological samples. BioRXiv doi: https://doi.org/10.1101/457820.

Guehrs, E., Schneider, M., Günther, C. M., Hessing, P., Heitz, K., Wittke, D., López-Serrano Oliver, A., Jakubowski, N., Plendl, J., Eisebitt, S., & Haase, A. (2017). Quantification of silver nanoparticle uptake and distribution within individual human macrophages by FIB/SEM slice and view. Journal of Nanobiotechnology, 15(1), 21. doi.org/10.1186/s12951-017-0255-8

de Haan, K., Ballard, Z. S., Rivenson, Y., Wu, Y., & Ozcan, A. (2019). Resolution enhancement in scanning electron microscopy using deep learning. Scientific Reports, 9(1), 12050. doi.org/10.1038/s41598-019-48444-2

Haueter, S., Kawasumi, M., Asner, I., Brykczynska, U., Cinelli, P., Moisyadi, S., Bürki, K., Peters, A. H. F. M., & Pelczar, P. (2010). Genetic vasectomy-Overexpression of Prm1-EGFP fusion protein in elongating spermatids causes dominant male sterility in mice. Genesis, NA–NA. doi.org/10.1002/dvg.20598

Ho, Elaine. (2022). rosalindfranklininstitute/quoll: v0.0.1 (v0.0.1). Zenodo.

Howard, J., & Gugger, S. (2020). Fastai: A Layered API for Deep Learning. Information, 11(2), 108. doi.org/10.3390/info11020108

Jacquemyn, J., Cascalho, A., & Goodchild, R. E. (2017). The ins and outs of endoplasmic reticulum-controlled lipid biosynthesis. EMBO Reports, 18(11), 1905–1921. doi.org/10.15252/embr.201643426

Joy DC, Joy CS. (1996). Low voltage scanning electron microscopy. Micron 27:247–263. doi:10.1016/0968-4328(96)00023-6

Kaseda, K., McAinsh, A. D., & Cross, R. A. (2012). Dual pathway spindle assembly increases both the speed and the fidelity of mitosis. Biology Open, 1(1), 12–18. https://doi.org/10.1242/bio.2011012

Khavnekar S, Vrbovská V, Zaoralová M, Kelley R, Beck F, Kotecha A, Plitzko J, Erdmann PS. (2022). Optimizing Cryo-FIB Lamellas for sub-5Å in situ Structural Biology. BioRXiv doi:10.1101/2022.06.16.496417

Klein S, Wachsmuth-Melm M, Winter SL, Kolovou A, Chlanda P. (2021). Cryo-correlative light and electron microscopy workflow for cryo-focused ion beam milled adherent cells. Methods Cell Biol 2021;162:273–302. doi:10.1016/bs.mcb.2020.12.009

Klumpe, S., Fung, H. K., Goetz, S. K., Zagoriy, I., Hampoelz, B., Zhang, X., Erdmann, P. S., Baumbach, J., Müller, C. W., Beck, M., Plitzko, J. M., & Mahamid, J. (2021). A modular platform for automated cryo-FIB workflows. ELife, 10. https://doi.org/10.7554/eLife.70506

Knott G, Marchman H, Wall D, Lich B. (2008). Serial Section Scanning Electron Microscopy of Adult Brain Tissue Using Focused Ion Beam Milling. Journal of Neuroscience 28:2959–2964. doi:10.1523/JNEUROSCI.3189-07.2008

Koho S, Tortarolo G, Castello M, Deguchi T, Diaspro A, Vicidomini G. (2019). Fourier ring correlation simplifies image restoration in fluorescence microscopy. Nature Communications 10:3103. doi:10.1038/s41467-019-11024-z

Kounatidis I, Stanifer ML, Phillips MA, Paul-Gilloteaux P, Heiligenstein X, Wang H, Okolo CA, Fish TM, Spink MC, Stuart DI, Davis I, Boulant S, Grimes JM, Dobbie IM, Harkiolaki M. (2020). 3D Correlative Cryo-Structured Illumination Fluorescence and Soft X-ray Microscopy Elucidates Reovirus Intracellular Release Pathway. Cell 182:515–530.e17. doi:10.1016/J.CELL.2020.05.051

Li, C. H., & Tam, P. K. S. (1998). An iterative algorithm for minimum cross entropy thresholding. Pattern Recognition Letters, 19(8), 771–776. doi.org/10.1016/S0167-8655(98)00057-9

Mahamid J, Schampers R, Persoon H, Hyman A a, Baumeister W, Plitzko JM. (2015). A focused ion beam milling and lift-out approach for site-specific preparation of frozen-hydrated lamellas from multicellular organisms. Journal of Structural Biology 192:262–269.

Martynowycz, M. W., Shiriaeva, A., Clabbers, M. T. B., Nicolas, W. J., Weaver, S. J., Hattne, J., & Gonen, T. (2022). A robust approach for MicroED sample preparation of lipidic cubic phase embedded membrane protein crystals. BioRxiv. https://doi.org/10.1101/2022.07.26.501628

Lowe, D. G. (2004). Distinctive Image Features from Scale-Invariant Keypoints. International Journal of Computer Vision, 60(2), 91–110. doi.org/10.1023/B:VISI.0000029664.99615.94

Reimer L, Tollkamp C. (1980). Measuring the backscattering coefficient and secondary electron yield inside a scanning electron microscope. Scanning 3:35–39. doi:10.1002/sca.4950030105

Ronneberger, O., Fischer, P., & Brox, T. (2015). U-Net: Convolutional Networks for Biomedical Image Segmentation (pp. 234–241). doi.org/10.1007/978-3-319-24574-4_28

Rueden CT, Schindelin J, Hiner MC, DeZonia BE, Walter AE, Arena ET, Eliceiri KW. (2017). ImageJ2: ImageJ for the next generation of scientific image data. BMC Bioinformatics 18:529. doi:10.1186/s12859-017-1934-z

Santuy A, Rodriguez JR, DeFelipe J, Merchan-Perez A. (2018). Volume electron microscopy of the distribution of synapses in the neuropil of the juvenile rat somatosensory cortex. Brain Structure and Function 223:77–90. doi:10.1007/s00429-017-1470-7

Schindelin, J., Arganda-Carreras, I., Frise, E., Kaynig, V., Longair, M., Pietzsch, T., Preibisch, S., Rueden, C., Saalfeld, S., Schmid, B., Tinevez, J.-Y., White, D. J., Hartenstein, V., Eliceiri, K., Tomancak, P., & Cardona, A. (2012). Fiji: an open-source platform for biological-image analysis. Nature Methods, 9(7), 676–682. doi.org/10.1038/nmeth.2019

Seiter, J., Muller, E., Blank, H., Gehrke, H., Marko, D., & Gerthsen, D. (2014). Backscattered electron SEM imaging of cells and determination of the information depth. Journal of Microscopy, 254(2), 75–83. doi.org/10.1111/jmi.12120

Schwartz, J., Jiang, Y., Wang, Y., Aiello, A., Bhattacharya, P., Yuan, H., Mi, Z., Bassim, N., & Hovden, R. (2019). Removing Stripes, Scratches, and Curtaining with Nonrecoverable Compressed Sensing. Microscopy and Microanalysis, 25(3), 705–710. doi.org/10.1017/S1431927619000254

Scher N, Rechav K, Paul-Gilloteaux P, Avinoam O. (2021). In situ fiducial markers for 3D correlative cryo-fluorescence and FIB/SEM imaging. iScience 24:102714. doi:10.1016/J.ISCI.2021.102714

Schertel A, Snaidero N, Han HM, Ruhwedel T, Laue M, Grabenbauer M, Möbius W. (2013). Cryo-FIB/SEM: Volume imaging of cellular ultrastructure in native frozen specimens. Journal of Structural Biology 184:355–360.

Seiler H. (1983). Secondary electron emission in the scanning electron microscope. Journal of Applied Physics 54:R1–R18. doi:10.1063/1.332840

Shen K, Pender CL, Bar-Ziv R, Zhang H, Wickham K, Willey E, Durieux J, Ahmad Q, Dillin A. (2022). Mitochondria as Cellular and Organismal Signaling Hubs. Annual Review of Cell and Developmental Biology 38. doi:10.1146/annurev-cellbio-120420-015303

Sofroniew, N., Lambert, T., Evans, K., Nunez-Iglesias, J., Bokota, G., Winston, P., Peña-Castellanos, G., Yamauchi, K., Bussonnier, M., Doncila Pop, D., Can Solak, A., Liu, Z., Wadhwa, P., Burt, A., Buckley, G., Sweet, A., Migas, L., Hilsenstein, V., Gaifas, L.,… McGovern, A. (2022). napari: a multi-dimensional image viewer for Python.

Sørensen, T. (1948). A method of establishing groups of equal amplitude in plant sociology based on similarity of species and its application to analyses of the vegetation on Danish commons. Kongelige Danske Videnskabernes Selskab, 5(4), 1–34.

Spehner D, Steyer AM, Bertinetti L, Orlov I, Benoit L, Pernet-Gallay K, Schertel A, Schultz P. (2020). Cryo-FIB/SEM as a promising tool for localizing proteins in 3D. Journal of Structural Biology 211:107528. doi:10.1016/j.jsb.2020.107528

Thompson RF, Walker M, Siebert CA, Muench SP, Ranson NA. (2016). An introduction to sample preparation and imaging by cryo-electron microscopy for structural biology. Methods 100:3–15.

de Tombe, P. P., & ter Keurs, H. E. D. J. (2016). Cardiac muscle mechanics: Sarcomere length matters. Journal of Molecular and Cellular Cardiology, 91, 148–150. doi.org/10.1016/j.yjmcc.2015.12.006

Tucker JD, Siebert CA, Escalante M, Adams PG, Olsen JD, Otto C, Stokes DL, Hunter CN. (2010). Membrane invagination in Rhodobacter sphaeroides is initiated at curved regions of the cytoplasmic membrane, then forms both budded and fully detached spherical vesicles. Molecular Microbiology 76:833–847. doi:10.1111/j.1365-2958.2010.07153.x

Valtschanoff JG, Weinberg RJ. (2001). Laminar Organization of the NMDA Receptor Complex within the Postsynaptic Density. The Journal of Neuroscience 21:1211–1217. doi:10.1523/JNEUROSCI.21-04-01211.2001

Vitiello E, Moreau P, Nunes V, Mettouchi A, Maiato H, Ferreira JG, Wang I, Balland M. (2019). Acto-myosin force organization modulates centriole separation and PLK4 recruitment to ensure centriole fidelity. Nature Communications 10:52. doi:10.1038/s41467-018-07965-6

Watson ML. (1958). Staining of Tissue Sections for Electron Microscopy with Heavy Metals. The Journal of Biophysical and Biochemical Cytology 4:475–478. doi:10.1083/jcb.4.4.475

Zhu Y, Sun D, Schertel A, Martin-Fernandez ML, Freyberg Z, Zhang Correspondence P, Ning J, Fu X, Gwo PP, Watson AM, Zanetti-Domingues LC, Zhang P. (2021). Serial cryo-FIB/SEM Reveals Cytoarchitectural Disruptions in Leigh Syndrome Patient Cells ll. doi:10.1016/j.str.2020.10.003

